# Identification of a novel regulator of *Clostridioides difficile* cortex formation

**DOI:** 10.1101/2021.03.03.433760

**Authors:** Megan H. Touchette, Hector Benito de la Puebla, Carolina Alves Feliciano, Benjamin Tanenbaum, Monica Schenone, Steven A. Carr, Aimee Shen

**Affiliations:** Department of Molecular Biology and Microbiology, Tufts University School of Medicine, Boston, Massachusetts, USA; Broad Institute of MIT and Harvard, Cambridge, Massachusetts, USA.

**Author notes:** Address correspondence to Aimee Shen. These authors contributed equally to this work (Megan H. Touchette, Hector Benito de la Puebla). University of Massachusetts Medical School, Worcester, MA.

## Abstract

*Clostridioides difficile* is a leading cause of healthcare-associated infections worldwide. *C. difficile* infections are transmitted by its metabolically dormant, aerotolerant spore form. Functional spore formation depends on the assembly of two protective layers: a thick layer of modified peptidoglycan known as the cortex layer and a multilayered proteinaceous meshwork known as the coat. We previously identified two spore morphogenetic proteins, SpoIVA and SipL, that are essential for recruiting coat proteins to the developing forespore and making functional spores. While SpoIVA and SipL directly interact, the identities of the proteins they recruit to the forespore remained unknown. We used mass spectrometry-based affinity proteomics to identify proteins that interact with the SpoIVA-SipL complex. These analyses identified the Peptostreptococcaceae family-specific, sporulation-induced bitopic membrane protein CD3457 (renamed SpoVQ) as a protein that interacts with SipL and SpoIVA. Loss of SpoVQ decreased heat-resistant spore formation by ∼5-fold and reduced cortex thickness∼2-fold; the thinner cortex layer of Δ*spoVQ* spores correlated with higher levels of spontaneous germination (i.e., in the absence of germinant). Notably, loss of SpoVQ in either *spoIVA* or *sipL* mutants prevented cortex synthesis altogether and greatly impaired the localization of a SipL-mCherry fusion protein around the forespore. Thus, SpoVQ is a novel regulator of *C. difficile* cortex synthesis that appears to link cortex and coat formation. The identification of SpoVQ as a spore morphogenetic protein further highlights how Peptostreptococcaceae family-specific mechanisms control spore formation in *C. difficile*.

**Importance:** The Centers for Disease Control has designated *Clostridioides difficile* as an urgent threat because of its intrinsic antibiotic resistance. *C. difficile* persists in the presence of antibiotics in part because it makes metabolically dormant spores. While recent work has shown that preventing the formation of infectious spores can reduce *C. difficile* disease recurrence, more selective anti-sporulation therapies are needed. The identification of spore morphogenetic factors specific to *C. difficile* would facilitate the development of such therapies. In this study, we identified SpoVQ (CD3457) as a spore morphogenetic protein specific to the Peptostreptococcaceae family that regulates the formation of *C. difficile’s* protective spore cortex layer. SpoVQ acts in concert with the known spore coat morphogenetic factors, SpoIVA and SipL, to link formation of the protective coat and cortex layers. These data reveal a novel pathway that could be targeted to prevent the formation of infectious *C. difficile* spores.

## Introduction

The bacterial pathogen *Clostridioides difficile* is a leading cause of healthcare-associated infections worldwide. In the United States, it is responsible for an estimated ∼470,000 infections with an associated treatment cost of >$1 billion annually (1). *C. difficile* causes colitis and can lead to severe complications such as pseudomembranous colitis, toxic megacolon, and death, particularly in elderly populations (2). These complications are more frequent in cases of disease recurrence (3), which occurs in ∼20% of *C. difficile* infections (4).

Prior antibiotic usage predisposes individuals to *C. difficile* infection because antibiotics disrupt the endogenous microbiota that typically provides colonization resistance against *C. difficile* infection (5, 6). *C. difficile* can persist in the face of antibiotic treatment in part because its metabolically dormant spore form is inert to antibiotics. This metabolic dormancy allows *C. difficile* spores to survive in the presence of oxygen and transmit disease (7, 8). Accordingly, preventing spore formation can reduce *C. difficile* disease recurrence in mice (9).

The basic steps of *C. difficile* spore formation resemble those first defined in *Bacillus subtilis* (10, 11). Once sporulation is activated at the transcriptional level, the first morphological event of sporulation is the formation of a polar septum during asymmetric division. This event creates a larger mother cell and smaller forespore cell, both of which induce distinct transcriptional programs that drive specific morphological changes. The larger mother cell will engulf the smaller forespore in a process analogous to phagocytosis. Following engulfment, the forespore is surrounded by (i) an outer forespore membrane derived from the mother cell and (ii) an inner forespore membrane derived from the forespore. The mother cell then produces an extensive array of coat proteins that localize to and encase the forespore to form the concentric proteinaceous shells that make up the coat. This coat layer protects spores against oxidative, chemical, and enzymatic insults. The mother cell and the forespore also collaborate to synthesize a thick layer of modified peptidoglycan known as the cortex. This massive cell wall layer constrains the spore’s size and ultimately prevents water from entering the developing forespore even as the forespore imports the spore-specific small molecule, calcium dipicolinic acid. The low water content of the spore cytosol, also known as the core, impairs metabolic processes and is essential for maintaining metabolic dormancy (12, 13).

While the general morphological stages of spore formation are conserved between *C. difficile*, *B. subtilis*, and other mono-spore formers (14), the mechanisms used by these organisms to complete each morphological stage can differ significantly. For example, several spore morphogenetic proteins identified in *B. subtilis* have different functional requirements in *C. difficile* (10). These differences are particularly acute at the level of coat assembly. SpoVM is conserved across most spore formers (15) and is essential for both coat assembly around the *B. subtilis* forespore and cortex synthesis (16, 17). However, SpoVM is mostly dispensable for functional spore formation in *C. difficile*, although ∼30% of sporulating *C. difficile spoVM* mutant cells exhibit cortex abnormalities (18). *C. difficile* SpoIVA phenocopies *B. subtilis* SpoIVA in that both are strictly required for functional spore formation and coat encasement (19–21). However, unlike *B. subtilis* SpoIVA, *C. difficile* SpoIVA is not strictly needed for cortex synthesis (20, 21).

*B. subtilis* SpoVM and SpoIVA are both essential for cortex formation because defects in their localization around the forespore triggers a quality control pathway that leads to mother cell lysis (17). While this quality control pathway is absent in the Clostridia (22), SpoVM and SpoIVA nevertheless impact cortex synthesis in *C. difficile* because mutants lacking either of these proteins generate forespores with cortex abnormalities (18, 23).

Beyond these conserved spore morphogenetic proteins, *C. difficile* uses clostridial- and Peptostreptococcaceae family-specific spore morphogenetic proteins to mediate spore assembly. SipL is a clostridial-specific spore morphogenetic protein that directly binds SpoIVA (<u>S</u>poIVA Interacting <u>P</u>rotein L) and is required for other coat proteins to localize to and encase the forespore (20). Both these proteins are made early in sporulation under the control of the mother cell-specific sigma factor, σ^E^, the first sporulation-specific sigma factor that gets activated in the mother cell (24). A *C. difficile sipL* mutant resembles a *C. difficile spoIVA* mutant in that both fail to polymerize coat layers around the forespore and produce cortex layers that are often irregular in shape and thickness (18, 20, 25). CotL is a Peptostreptococcaceae family-specific coat morphogenetic protein that also regulates cortex thickness (26). Loss of CotL strongly impairs the localization and/or retention of coat and cortex-localized proteins in mature spores, which also produce a thinner cortex layer. Taken together, these analyses suggest an intriguing but poorly understood link between coat and cortex assembly in *C. difficile*. In this study, we identify SpoVQ as a novel Peptostreptococcaceae-family protein that appears to link these two morphological processes. In particular, SpoVQ regulates cortex synthesis and genetically and physically interacts with the SpoIVA and SipL coat morphogenetic proteins.

### Identification of SipL-binding proteins using co-immunoprecipitation

Both SpoIVA and SipL are landmark proteins for coat morphogenesis. Loss of either of these proteins results in polymerized coat mislocalizing to the cytosol or failing to fully encase the forespore (20, 23, 25). Our previous work suggests that SpoIVA and SipL are recruited as a complex to the forespore (25) and that binding between these two proteins facilitates their encasement of the forespore (23). To identify interacting partners of SipL and potentially SpoIVA, we immunoprecipitated members of the complex and analyzed the pulldowns by quantitative mass spectrometry-based proteomics (27, 28). Specifically, we immunoprecipitated a FLAG-tagged SipL variant that we previously showed pulls down untagged SpoIVA (23). The strain producing this variant expresses a *sipL* construct encoding SipL with three tandem FLAG tags at its C-terminus from the ectopic *pyrE* locus of a Δ*sipL* mutant strain (25). The *pyrE* locus was used for all complementation strains generated in this study (29)). To determine whether proteins beyond SpoIVA could be identified in SipL-FLAG_3_ pull-downs, we Coomassie-stained FLAG peptide elutions from co-immunoprecipitations with the *ΔsipL/sipL-FLAG_3_* strain orthe untagged Δ*sipL* complementation strain. Several bands representing potential interacting partners were observed in the elution fraction of the SipL-FLAG_3_ coimmunoprecipitation but not in the untagged SipL control sample (**Figure 1A**). The band at 65 kDa is likely SpoIVA based on western blot analyses (**Figure 1B**).

**Figure 1.**
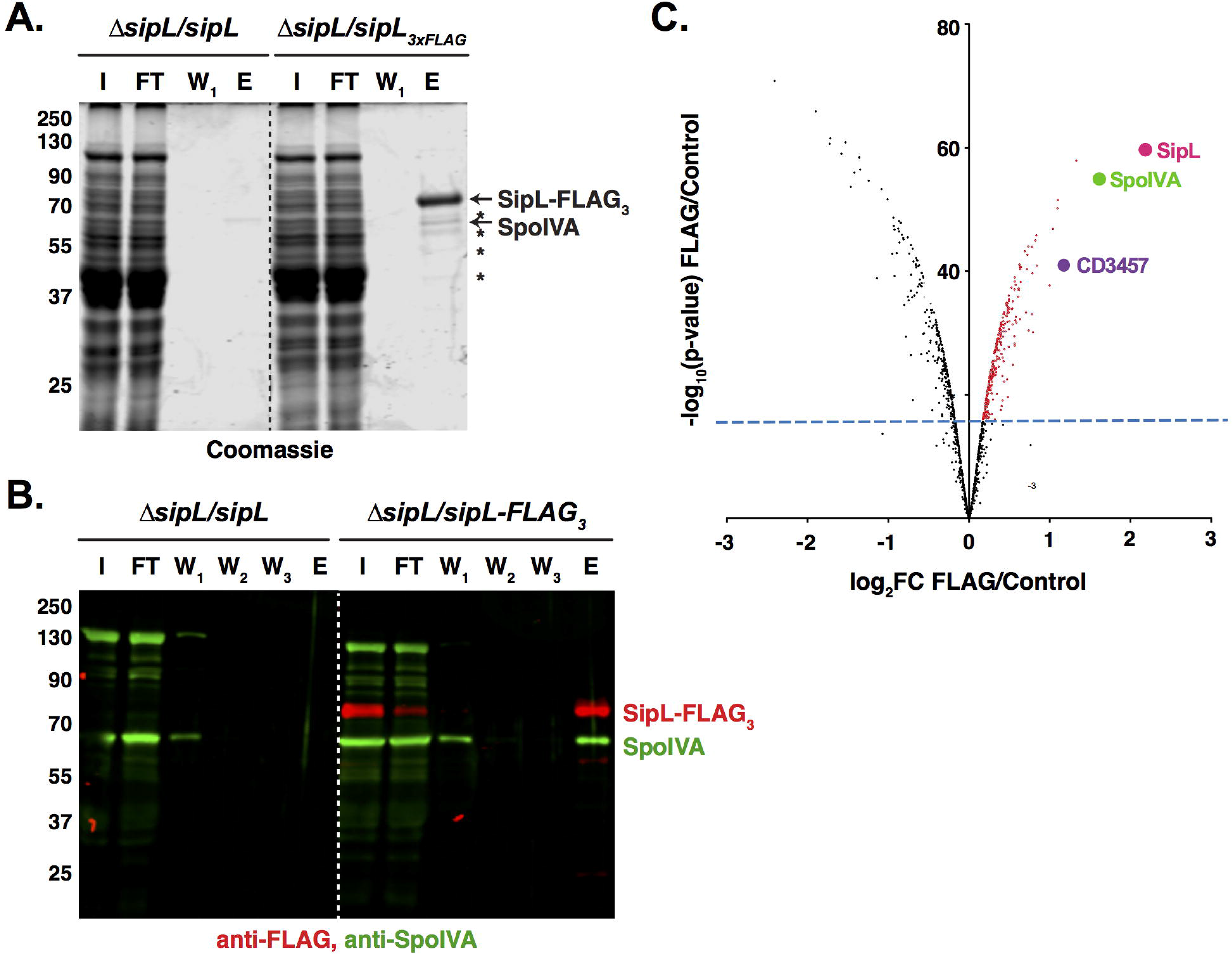
Mass spectrometry analyses of SipL co-immunoprecipitations identify SpoVQ as an interacting partner. (A) Coomassie stain of SipL co-immunoprecipitation fractions reveals potential interacting partners highlighted by asterisks. I = Input, FT = flowthrough, W_1_ = first wash, W_2_ = second wash, W_3_ = third wash, E = FLAG-peptide elution. (B) Western blot analysis detecting SpoIVA and SipL in SipL co-immunoprecipitation fractions. SipL-FLAG_3_ was detected using an anti-FLAG antibody (red), while untagged SpoIVA was detected using an anti-SpoIVA antibody (green). I = Input, FT = flowthrough, W_1_ = first wash, W_2_ = second wash, W_3_ = third wash, E = FLAG-peptide elution. (C) Volcano plot of mass spectrometry data showing enrichment of SipL interacting proteins in red that exceeded the minimum statistical threshold of p < 0.01 with a fold-change (FC) ≥ 2.

To identify additional SipL-interacting proteins, we performed trypsin on-bead digestions of the co-immunoprecipitation samples and labeled the digestions with isobaric tag reagents. After equally mixing the labeled samples, the pooled sample was subjected to quantitative LC-MS/MS. Analyses of these samples confirmed SpoIVA as a SipL-interacting partner (3-fold enrichment, p < 10^-6^) and identified a number of cytosolic proteins (**Figure 1C**). Of the handful of proteins enriched >2-fold, one candidate, CD3457, stood out because (i) its gene is strongly induced in the mother cell under the control of σ^E^ during sporulation similar to *spoIVA* and *sipL* (30, 31); (ii) it was identified in transposon mutagenesis screen for sporulation mutants (32); and (iii) it is conserved exclusively in the Peptostreptococcaceae family.

### CD3457 (SpoVQ) is a transmembrane protein that regulates heat-resistant spore formation

To determine whether CD3457 (renamed SpoVQ for reasons detailed below) regulates coat formation and/or functional spore formation in *C. difficile*, we deleted Δ*spoVQ* from *C.difficile630*Δ*erm*Δ*pyrE* (29) and analyzed the sporulation phenotype of the resulting mutant *difficile* using a heat resistance assay. This assay measures the capacity of sporulating cultures to produce spores that can survive a heat treatment that kills vegetative cells and germinate on media containing bile acid germinant (33). Loss of *spoVQ* resulted in a 5-fold reduction in heat-resistant spore formation (p < 0.0001). This defect was largely complemented by expressing a wild-type copy of *spoVQ* from the ectopic *pyrE* locus (**Figure 2A**). Western blot analyses Δ*spoVQ*/*spoVQ* complementation strain produces SpoVQ at wild-type levels (**Figure 2B**) and indicated that SpoIVA and SipL levels were unaffected by loss of SpoVQ.

**Figure 2.**
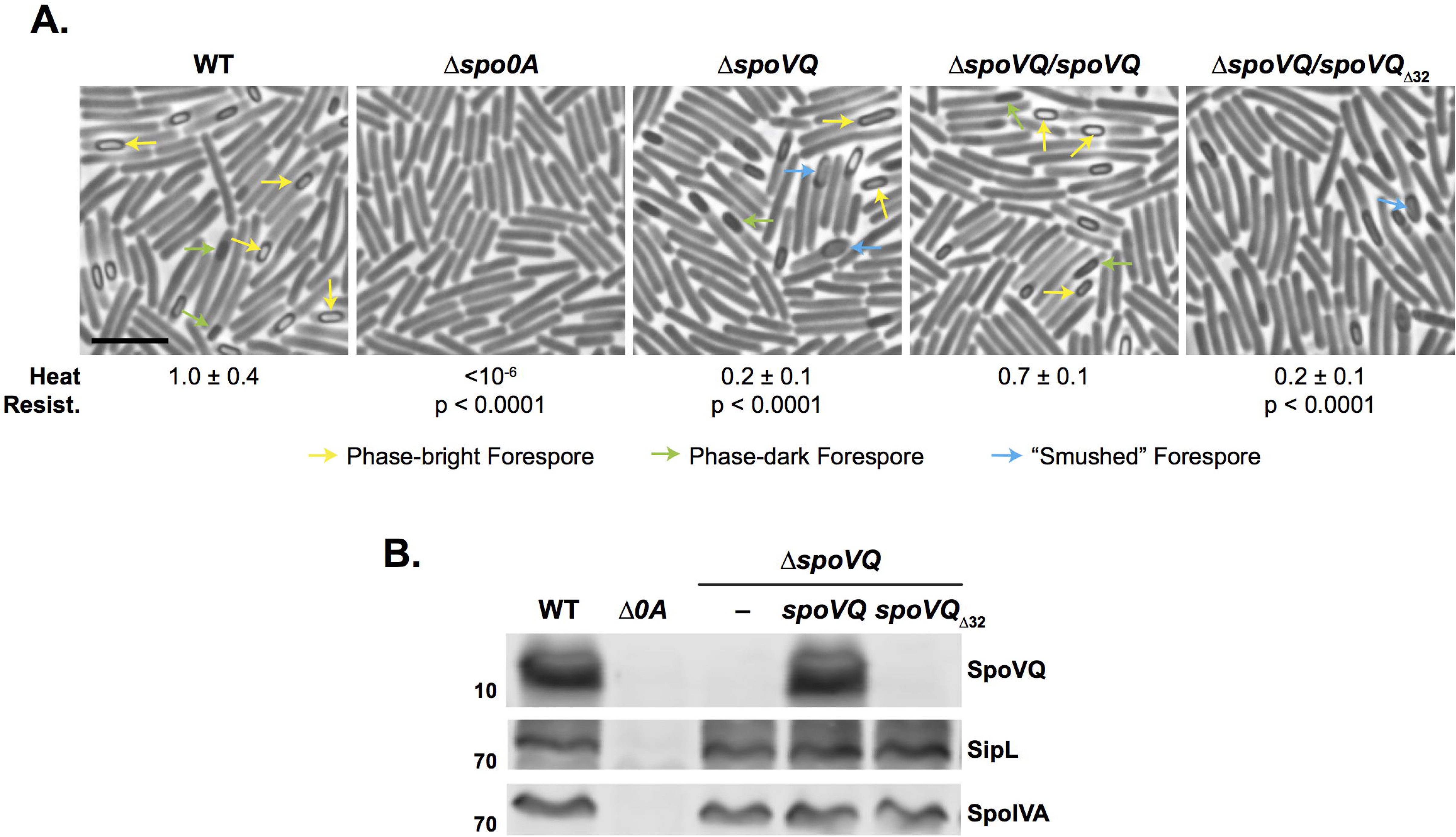
Loss of SpoVQ results in morphological and functional defects in spore formation. (A) Phase-contrast microscopy images of sporulating cultures of wild-type 630Δ*erm*-p (WT, *pyrE* restored) and the indicated strains ∼20 hrs after sporulation induction. Δ*spoVQ* was complemented with either wild-type *spoVQ* or *spoVQ* _Δ32_, the latter of which encodes an N-terminal truncation of SpoVQ’s transmembrane domain. Arrows mark mature phase-bright forepores (yellow), immature phase-dark forespores (green), and “smushed” forespores (blue), which appeared flattened relative to the oblong phase-dark or phase-bright forespores highlighted. Heat resistance efficiencies were calculated from 20-24 hr sporulating cultures. The efficiencies represent the average ratio of heat-resistant spore CFUs to total cells for a given strain relative to WT based on a minimum of three biological replicates. The limit of detection of the assay is 10^-6^. Scale bar represents 5 µm. (B) Western blot analyses of strains shown in (A) using anti-SpoVQ, anti-SipL (20), and anti-SpoIVA antibodies.

Although SpoVQ is annotated as a conserved hypothetical protein, domain analyses predicted that SpoVQ carries an N-terminal transmembrane domain (**Figure S1A**). To confirm that SpoVQ carries an N-terminal transmembrane domain, we compared the solubility of recombinant full-length SpoVQ relative to a variant lacking the N-terminal 32 residues in *Escherichia coli*. Consistent with the N-terminal 32 aa forming a transmembrane domain (SpoVQ_Δ32_), full-length SpoVQ was primarily observed in the insoluble fraction, whereas SpoVQ _Δ32_ was highly soluble (**Figure S1B**).

To test whether the putative transmembrane domain is important for SpoVQ function, we complemented our Δ*spoVQ* strain with a construct that deletes the region encoding the transmembrane domain (*spoVQ*_Δ32_). This truncation construct failed to complement the heat resistance defect of Δ*spoVQ* (**Figure 2A**). This functional defect likely results from the truncation destabilizing SpoVQ, since SpoVQ_Δ32_ was undetectable in western blot analyses of sporulating cell lysates (**Figure 2B**) even though the anti-SpoVQ antibody used was raised against SpoVQ_Δ32_.

To determine what stage of sporulation was being impacted by loss of SpoVQ, we Δ*spoVQ* mutant strains by phase-contrast microscopy. Sporulating cultures of Δ*spoVQ* appeared to sporulate with similar frequency as wild type, but fewer mature phase-Δ*spoVQ* relative to wild type (**Figure 2A**, yellow arrows). Δ*spoIVA* and sporulating cells (20, 25, 34), mislocalized coat was not seen in Δ*spoVQ* sporulating cells. However, Δ*spoVQ* cells frequently produced phase-dark forespores with a “smushed” appearance, a phenotype we had not previously observed in *C. difficile* mutants (**Figure 2A**, blue arrows). This “smushed” forespore phenotype was reversed in wild-type *spoVQ* but not the Δ*spoVQ*_Δ32_ complementation strains, consistent with the heat-resistance phenotypes measured.

### Cortex formation is impaired in Δ*spoVQ* spores

The “smushed” appearance of the spores led us to examine whether Δ*spoVQ* spores would be more difficult to purify. To test end, we compared the spore purification efficiencies of Δ*spoVQ/spoVQ* strains. Loss of SpoVQ resulted in a ∼3-fold decrease in spore purification efficiency across three biological replicates (p < 0.0005, **Figure 3A**), a decrease that was largely reversed in the Δ*spoVQ*/*spoVQ* complementation strain. The Δ*spoVQ* spores that survived the purification procedure were largely indistinguishable from wild-type spores when analyzed by phase-contrast microscopy (**Figure S2**), although they were often less phase-bright (blue arrows, **Figure S2**). Consistent with previous observations that spore refractivity correlates with cortex synthesis (12), *spoVQ* mutant spores in transmission electron microscopy (TEM) analyses produced cortex layers that were ∼60% the thickness of wild-type spores (66 ± 13 nm vs. 38 ± 9 nm, p < 0.0001, **Figure 3B, 3C**). The cortex layer of the Δ*spoVQ*/*spoVQ* complementation strain was ∼80% the thickness of wild-type spore (p < 0.0001, **Figure 3B**), but the mean cortex thickness was still significantly higher than that of the parental Δ*spoVQ* mutant (p < 0.0001). Finally, the coat layers of Δ*spoVQ* spores were similar in thickness and appearance to wild-type spores, suggesting that loss of SpoVQ specifically affects cortex thickness (**Figure 3C**).

Interestingly, the thinner cortex of Δ*spoVQ* spores resembled that of *cotL* mutant spores (26). Like SpoVQ, CotL is a mother cell-specific, σ^E^-regulated protein found exclusively in the Peptostreptococcaceae family that affects cortex thickness (26). Loss of CotL also causes coat encasement defects and reduced incorporation of proteins predicted to be part of the cortex layer. To compare the *spoVQ* and *cotL* mutant phenotypes more directly, we analyzed the levels of coat and putative cortex proteins in Δ*spoVQ* and *cotL* mutant spores. These western blot analyses indicated that Δ*spoVQ* spore contain wild-type levels of outer coat proteins like CotA (35) and putative cortex-localized proteins like the SleC cortex lytic enzyme and germinant signaling proteins, CspC and CspB (10) (**Figure S2B**). In contrast Δ*cotL* spores contained dramatically reduced levels of these proteins (and of the basement layer proteins, SpoIVA and SipL, (25)) similar to prior analyses of a *cotL* targetron mutant (26). Thus, while the cortex thickness phenotype of Δ*spoVQ* spores resembled those of Δ*cotL* spores, these two mutant strains exhibit notable differences in their protein composition (**Figure S2B**), suggesting that they impact *C. difficile* spore formation using different mechanisms.

**Figure 3.**
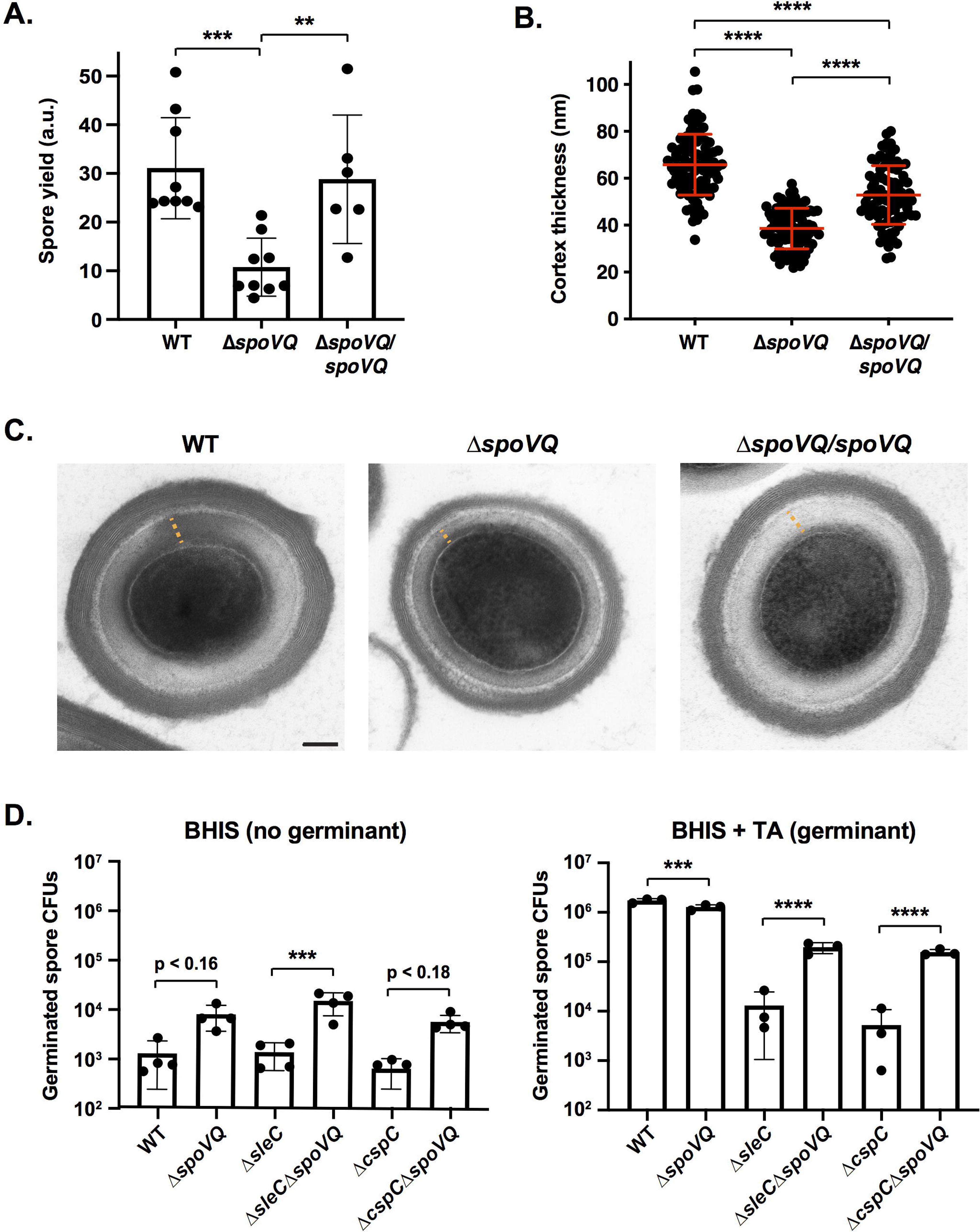
Δ*spoVQ* spores are purified less efficiently, produce a thinner cortex layer, andare prone to spontaneous germination. (A) Spore yields from purifications of WT, Δ*spoVQ* and Δ*spoVQ* complementation strains from a minimum of six biological replicates. Yields were determined by measuring the optical density of spore purifications at 600 nm and are expressed in arbitrary units (a.u.). Statistical significance relative to WT was determined using one-way ANOVA and Tukey’s test. *** p < 0.0005, ** p < 0.01. (B) Cortex thickness (nm) of the indicated spores based on transmission electron microscopy (TEM) analyses. Measurements were made on a minimum of 85 spores per strain and are representative of two biological replicates. **** p < 0.0001. (C) TEM of the indicated spores. Orange dashed line highlights the cortex thickness measured. Scale bar represents 100 nm. (D) Spontaneous germination of WT, Δ*spoVQ*, Δ*sleC* and Δ*cspC* germination mutants and the indicated double mutants spores on rich (BHIS) media lacking germinant (BHIS alone) and rich media containing germinant (BHIS + taurocholate (TA)). Germination data is based on analyses of four independent spore preparations. Statistical significance relative to the parental strain is shown and derives from one-way ANOVA analyses and Tukey’s test. **** p < 0.0001, *** p < 0.005.

### *spoVQ* mutant spores are prone to spontaneous germination

The thinner cortex layer of Δ*spoVQ* spores prompted us to consider whether these mutant spores would be more likely to prematurely germinate. This question was prompted by the “smushed” morphology of Δ*spoVQ* forespores, which somewhat resembled the phase-dark appearance of *Bacillus subtilis* mutant spores that prematurely germinate due to defects in spore assembly (36). In these *B. subtilis* mutants, deletion of germinant receptor genes partially rescues their heat-resistant spore formation defects, many of which are due to impaired cortex formation. Δ*spoVQ* forespores with a “smushed” appearance were prematurely germinating, we assessed whether loss of key germinant signaling proteins would prevent premature Δ*spoVQ* spores. For these analyses, we deleted *cspC*, which encodes the likely bile acid germinant receptor (37, 38), or *sleC*, which encodes the cortex lytic enzyme that degrades the protective cortex layer during germination (39, 40). These proteins are critical for bile acid germinant signal transduction and thus *C. difficile* spore germination (37, 39, 40).

However, contrary to our hypothesis, deletion of *sleC* or *cspC* did not improve the purification efficiency of Δ*spoVQ* spores (data not shown). In fact, the Δ*spoVQ*Δ*sleC* and Δ*spoVQ*Δ*cspC* double mutants germinated at higher levels on rich media containing germinant than single mutants lacking either of these critical germination proteins by 15-to 30-fold, respectively (**Figure 3D**); this enhanced germination was not statistically significant relative to wild type.

Notably, while Δ*spoVQ* spores germinated at close to wild-type levels on rich media containing germinant (1.4-fold reduction, p < 0.001), when the mutant spores were plated on rich media *lacking* germinant, they germinated at ∼6-fold higher levels than wild-type spores (p ∼ 0.17). Furthermore, deletion of spoVQ from either Δ*sleC* or Δ*cspC* Δ*cspC* spores enhanced this spontaneous germination in the absence of germinant by 14- and 10-fold, respectively (**Figure 3D**); the former result was statistically significant, p < 0.0005; the latter was not. Since spontaneous germination has been defined as spore germination that occurs in the absence of (i) germinant and/or (ii) key germinant signaling proteins like germinant receptors (e.g. CspC) or cortex lytic enzymes (e.g. SleC) (41–43), our results indicate that loss of SpoVQ promotes spontaneous germination of wild-type and germinant signaling mutant spores, a phenotype that could be linked to the thinner cortex of Δ*spoVQ* spores (**Figure 3B**).

### SpoVQ is required for cortex formation in the absence of SpoIVA or SipL

Since SpoVQ affects cortex synthesis (**Figure 3**) and binds SipL and/or SpoIVA (**Figure 1**), which themselves impact cortex thickness (23, 25), we wondered whether SpoVQ was required for cortex synthesis in Δ*spoIVA* or Δ*sipL* strains. To test this possibility, we deleted *spoVQ* from either Δ*spoIVA* or Δ*sipL* strains and analyzed the resulting double mutants morphologically using phase-contrast microscopy and TEM analyses. While developing forespores with phase-dark outlines were observed in WT, Δ*spoIVA*, Δ*sipL*, and Δ*spoVQ* Δ*spoVQ* strains (orange arrows), these dark outlines were not observed in the double mutant Δ*spoIVA*Δ*spoVQ* and Δ*sipL*Δ*spoVQ* strains (**Figure S3A**). Phase-dark outlines around forespores are typically observed when forespores become phase-bright due to the dehydration of the forespore cytosol as it matures (12), which occurs when the thick cortex layer is synthesized. To assess whether Δ*spoIVA*Δ*spoVQ* and Δ*sipL*Δ*spoVQ* strains produced a cortex layer, we analyzed sporulating cultures of these cells using TEM.

In developing forespores, mature cortex appears thick and electron-light (white) in the micrographs due to the large amount of cortex peptidoglycan synthesized during sporulation and its reduced level of crosslinking (12). In contrast, immature cortex layers are thinner and darker (dark gray) in electron micrographs. Wild-type forespores primarily produced a thick cortex layer that was electron-light (white), i.e. fully mature (**Figures 4A** and **4B**). A thick, mature cortex layer was also observed in ≈30% of Δ*spoIVA* and Δ*sipL* forespores, consistent with our prior report that SpoIVA and SipL are not essential for cortex formation in *C. difficile* (20). Nevertheless, as described earlier, Δ*spoIVA* and Δ*sipL* forespores exhibit cortex abnormalities, Δ*sipL* sporulating cells making cortex that is thinner and/or Δ*sipL* sporulating cells frequently exhibited indentations or areas of abnormal thickness as previously described (18, 23, 25). In contrast, Δ*spoVQ* sporulating cells rarely produced a thick, mature cortex layer (∼10%, blue arrows, **Figures 4B** and **S3B**); The cortex in Δ*spoIVA* and Δ*sipL* sporulating cells instead, this layer was markedly darker and thinner (**Figure 4A**). Notably, the cortex layer was even thinner Δ*spoIVA*Δ*spoVQ* and Δ*sipL* Δ*spoVQ* sporulating cells, with the majority of cells in the double mutants producing a dark, thin cortex layer that was difficult to distinguish from the germ cell wall. The double mutants, like the parental mutants Δ*sipL* and Δ*spoIVA*, frequently produced forespores with abnormal shapes. Taken together, these results strongly suggest that formation of the cortex layer in Δ*spoIVA* and Δ*sipL* mutants depends on the presence of SpoVQ, further highlighting the importance of this hypothetical protein in regulating cortex formation.

**Figure 4.**
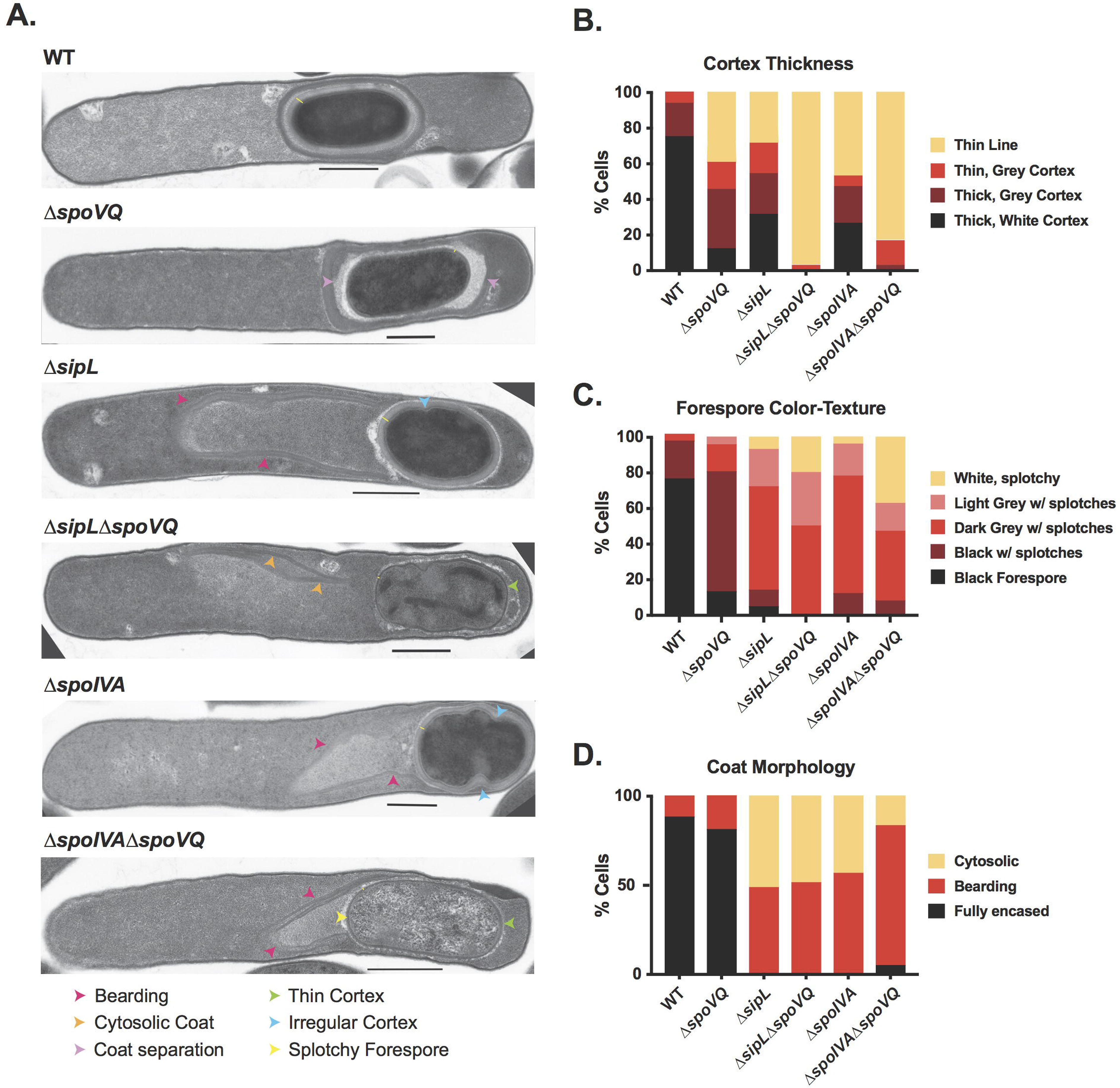
SpoVQ regulates cortex formation in a SpoIVA- and SipL-dependent manner. (A) Transmission electron microscopy (TEM) of wild-type (WT), Δ*spoVQ*, Δ*sipL*, Δ*spoIVA*, and the indicated double mutants 23 hrs after sporulation induction. Arrows highlight various phenotypes observed in the different mutant strains. Δ*spoVQ* sporulating cells exhibited many of these phenotypes. (B) Cortex thickness and electron density in the indicated strains. Mature cortex appears as an electron-light layer on top of a darker germ cell wall. Cortex designated “Thin line” likely represents the darker germ cell wall of the forespore, a phenotype that was particularly prominent in Δ*spoIVA*Δ*spoVQ* and Δ*sipLΔspoVQ* mutants (green arrow in (A)). (C) Electron density of forespores in the indicated strains as described by color and texture. Electron-dark, uniform forespores likely represent mature forespores whose core (cytosol) has partially dehydrate. Splotches likely arise during TEM sample processing and may be more likely in forespores that are more hydrated. (D) Coat morphology of the indicated strains. “Fully encased” represents polymerized coat encircling the forespore. “Bearding” refers to polymerized coat sloughing off the forespore (pink arrows in (A)). Cytosolic refers to polymerized coat mislocalizing entirely from the forespore into the cytosol (orange arrows in (A)).

The reduced thickness and higher electron density of the cortex layer in Δ*spoVQ* spores correlated with lighter electron density in the forespore cytosol (also known as the core, **Figures 4C** and **S3C**). Less than 20% of Δ*spoVQ* spores produced an electron-dense (dark) forespore, which could suggest that Δ*spoVQ* spores may be more hydrated than wild-type spores. The forespores of Δ*spoIVA* and Δ*sipL* derivatives were also less electron dense and thus inversely correlated with the thickness of their cortex layers.

Despite these differences in forespore core and cortex appearance in Δ*spoVQ* cells, coat layers appeared to encase Δ*spoVQ* forespores (**Figure 4D**), a finding that is consistent with our western blot studies of Δ*spoVQ* spores (**Figure S2**). As reported previously, loss of either SpoIVA or SipL largely abrogated coat encasement of the forespore irrespective of whether SpoVQ was present. In Δ*sipL* and Δ*spoIVA* mutants, polymerized coat was observed sloughing off the forespore (termed “bearding,”) or completely mislocalized to the cytosol (**Figure 4D**). Thus, despite SpoVQ’s effect on cortex thickness, it does not appear to affect coat encasement, again in contrast with CotL’s effect on both coat encasement and cortex formation in *C. difficile* (26).

### SpoVQ specifically localizes to forespore membranes in a SpoIVA- and SipL-independent manner, although SpoVQ affects SipL localization to the forespore

Our finding that SpoVQ regulates cortex synthesis in a SpoIVA- and SipL-dependent manner prompted us to consider whether SpoVQ affected SpoIVA and/or SipL localization around the forespore and vice versa. To address these questions, we analyzed the localization dependencies of SpoVQ, SpoIVA, and SipL using fluorescent protein fusions. We constructed a C-terminal mCherry fusion to SpoVQ and analyzed its localization around the forespore in merodiploid and Δ*spoVQ* backgrounds. In both these strain backgrounds, SpoVQ-mCherry specifically localized around the forespore (**Figure 5**). This observation suggests that even though SpoVQ-mCherry should have the capacity to localize to all mother cell-derived membranes, since it is produced under the control of σ^E^ (30, 31), it concentrates within the mother cell-derived forespore membrane through an unknown mechanism. Importantly, the SpoVQ-mCherry fusion fully complementedx Δ*spoVQ* (data not shown) and primarily produced a full-length fluorescent protein fusion (**Figure S4**).

**Figure 5.**
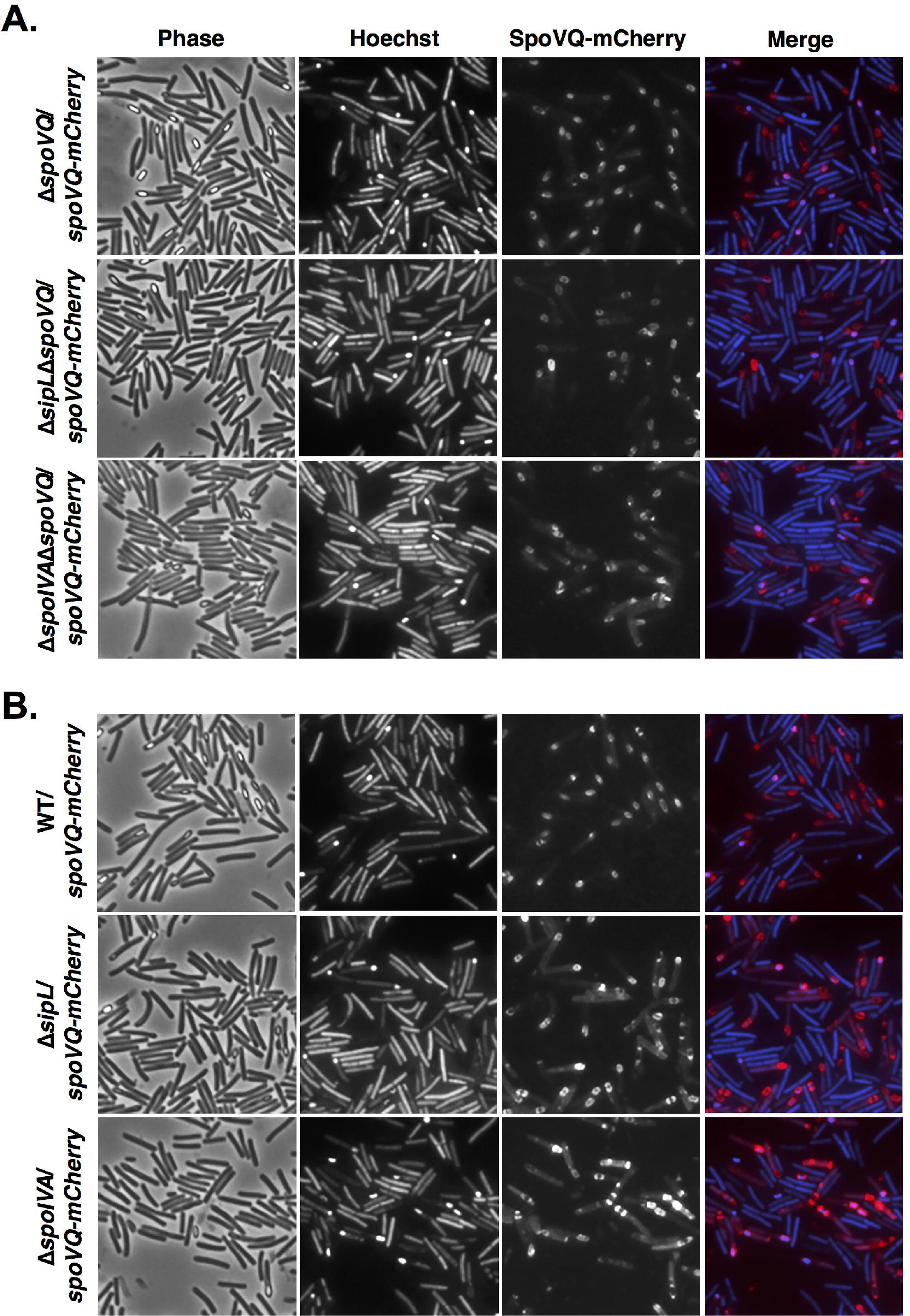
SpoVQ localizes around the forespore in a largely SpoIVA- and SipL-independent manner. (A) SpoVQ-mCherry localization in Δ*spoVQ*, Δ*sipL*Δ*spoVQ*, and Δ*spoIVA*Δ*spoVQ* strains, where the fusion protein is the only version of SpoVQ present. (B) SpoVQ-mCherry localization in wild type, Δ*sipL*, and Δ*spoIVA* strains, where the fusion protein and untagged SpoVQ are both produced. A slight increase in cytosolic signal in SpoVQ-mCherry was observed in the Δ*spoIVA* strain background. Sporulating cells were visualized by phase-contrast microscopy (Phase); the nucleoid was visualized with Hoechst. SpoVQ-mCherry fluorescence is shown in red, and Hoechst staining in blue in the “Merge” image. Images are representative of the results of three biological replicates.

Since our biochemical (**Figure 1**) and genetic data (**Figure 4**) suggested an interaction between SpoVQ and the SipL-SpoIVA complex, we assessed whether SpoIVA or SipL affects SpoVQ’s localization around the forespore. To this end, we complemented our Δ*spoIVA*, Δ*sipL*, *spoIVA*, Δ*sipL* Δ*spoVQ* double mutant strains with Δ*spoVQ-mCherry* and analyzed the localization of protein fusion during sporulation. SpoVQ-mCherry primarily localized around the forespore in the absence of either SpoIVA or SipL (**Figure 5**), regardless of whether untagged SpoVQ was also made. However, loss of SpoIVA seemed to increase the cytosolic SpoVQ-mCherry signal. These results suggest that SpoIVA may promote SpoVQ localization around the forespore, but neither SpoIVA nor SipL is absolutely essential for SpoVQ to concentrate in the mother cell-derived outer forespore membrane.

To determine whether SpoVQ affects SpoIVA and/or SipL localization around the forespore, we analyzed the localization of previously published mCherry-SpoIVA and SipL-mCherry fusions in the presence and absence of SpoVQ. To localize mCherry-SpoIVA, we introduced a construct encoding an mCherry-SpoIVA fusion protein (25) into the *pyrE* locus of Δ*spoVQ* mutant. It was necessary to generate this *spoIVA* merodiploid strain because untagged SpoIVA promotes encasement of the partially functional mCherry-SpoIVA fusion (18). SpoVQ did not appear to affect localization of mCherry-SpoIVA around the forespore (**Figure 6A**), indicating that SpoIVA forespore encasement does not depend on SpoVQ. In contrast, localization of a functional SipL-mCherry fusion protein was significantly impaired in the absence of SpoVQ, with much of the SipL-mCherry signal redistributing to the cytosol of Δ*sipL*Δ*spoVQ*/*sipL-mCherry* cells (**Figure 6B**). In this strain, SipL-mCherry is the only copy of SipL, since we previously showed that co-production of the fusion with untagged SipL increased the cytosolic signal of SipL-mCherry (25). Western blot analyses indicated that SipL-mCherry levels were elevated in the Δ*spoVQ* background relative to wild type and the Δ*sipL* complementation strain (**Figure S5**). The increased mCherry signal in the cytosol of edistributing to the cytosol of Δ*sipL*Δ*spoVQ*/*sipL-mCherry* could be due to increased liberation of mCherry from the higher levels of SipL-mCherry. Despite this degradation, some SipL-mCherry fusion was observed around the forespore (**Figure 6B**, yellow arrows), although the functionality of the fusion was reduced ≈ 10-fold relative toΔ*spoVQ* alone (**Figure S5**). Indeed, ∼100-fold fewer heat-resistant spores were detected in Δ*sipL*Δ*spoVQ/sipL-mCherry* relative to Δ*sipL/sipL-mCherry* relative to **S5**)

**Figure 6.**
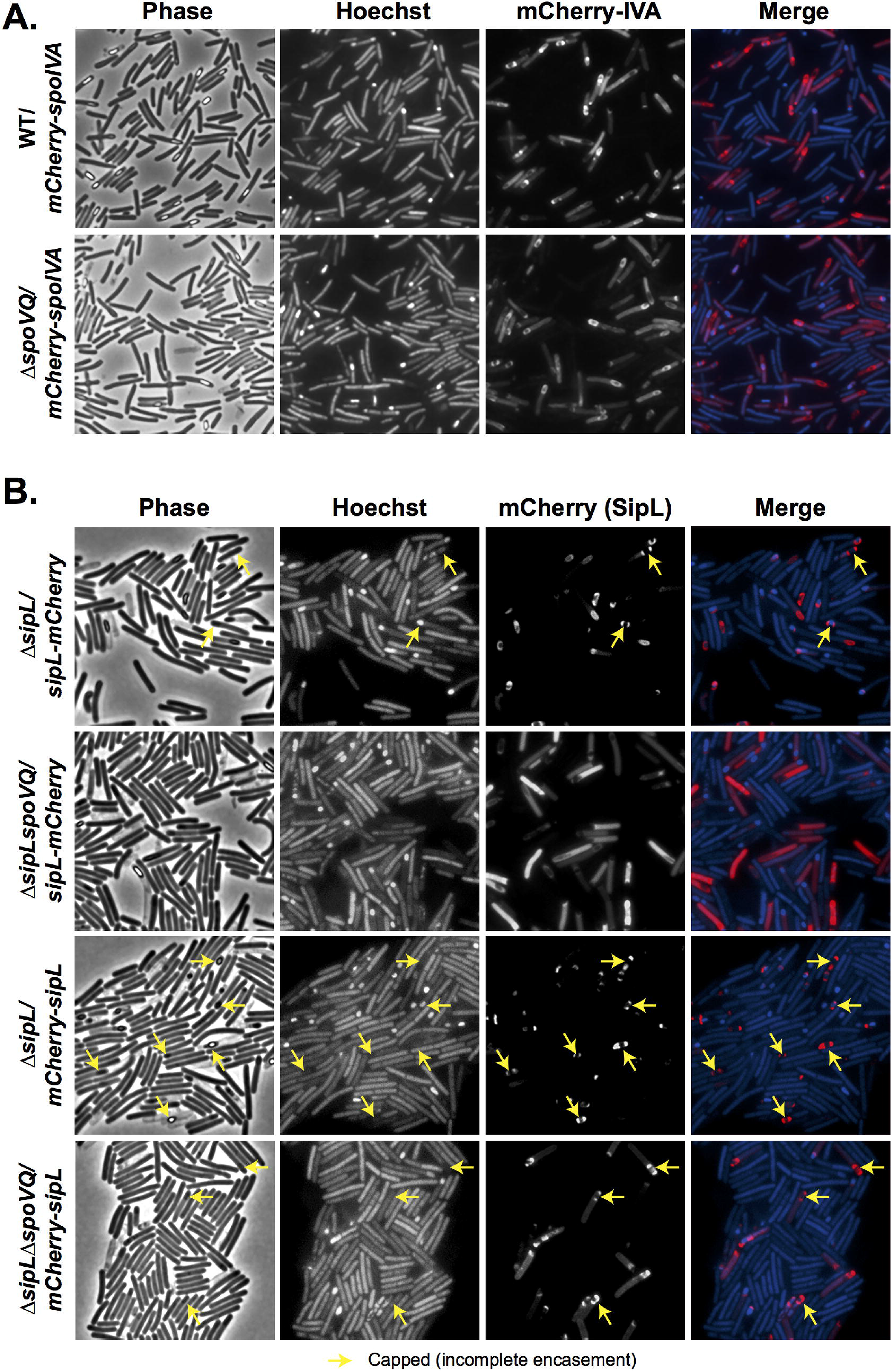
SpoVQ affects the localization of SipL-mCherry but not mCherry-SipL or mCherry-SpoIVA fusions. (A) mCherry-SpoIVA localization in wild-type versus Δ*spoVQ* strains, where the fusion protein and untagged SpoIVA are both produced. (B) SipL-mCherry or mCherry-SipL (C-terminal vs. N-terminal mCherry fusion, respectively) localization in Δ*sipL* versus *spoVQ* strain backgrounds. In these strains, the fusion protein is the only version of SipL present. The *mCherry-sipL* fusion construct complements Δ*sipL* less efficiently than Δ*sipL-mCherry* in terms of heat-resistant spore formation (**Figure S5**), consistent with mCherry-SipL encasing the forespore less efficiently than SipL-mCherry (yellow arrows, “capped” distribution). Cytosolic mCherry signal was elevated in the absence of SpoVQ, particularly with the C-terminal SipL-mCherry fusion. Sporulating cells were visualized by phase-contrast microscopy (Phase); the nucleoid was visualized with Hoechst. mCherry fluorescence of the indicated fusion proteins is shown in red, and Hoechst staining in blue in the “Merge” image. Images are representative of the results of three biological replicates.

Given that loss of SpoVQ caused much of the SipL-mCherry signal to mislocalize to the cytosol and reduced the function of this fusion protein by ∼10-fold (**Figure S5**), we wondered whether the fusion of mCherry to the C-terminus of SipL was interfering with SipL function specifically in the absence of SpoVQ. SipL’s C-terminal LysM domain binds directly to SpoIVA (20, 25) and may also bind peptidoglycan (25, 44, 45), so fusions to SipL’s C-terminus could disrupt binding to SpoVQ, if SpoVQ is oriented into the intermembrane space. Consistent with this hypothesis, moving the mCherry fusion to the N-terminus of SipL increased localization of the mCherry-SipL to the forespore even in the absence of SpoVQ (**Figure 6B**). In particular, the majority of the mCherry signal in Δ*sipL*Δ*spoVQ*/*mCherry-SipL* localized to the forespore in contrast with Δ*sipL*Δ*spoVQ*/*sipL-mCherry*. Nevertheless, loss of SpoVQ increased cytosolic mCherry-SipL signal in Δ*sipL*Δ*spoVQ* relative to the wild-type *sipL* complementation strain.

Surprisingly, the function of the N-terminal mCherry-SipL was reduced even further than the C-terminal fusion in the absence of SpoVQ (∼10-fold fewer heat-resistant spores were observed in Δ*sipL*Δ*spoVQ/mCherry-sipL* relative to Δ*sipL*Δ*spoVQ*/*sipL-mCherry*, **Figure S5**). The reduced function of mCherry-SipL may simply reflect the reduced levels of this fusion protein relative to SipL-mCherry or untagged SipL (**Figure S5**). This reduction in mCherry-SipL may explain why the mCherry-SipL fusion (i.e. Δ*sipL/mCherry-sipL*) encased the forespore less efficiently than SipL-mCherry (**Figure 6B**). The Δ*sipL/sipL-mCherry* foresporesalso appeared more round than wild-type and Δ*sipL/sipL-mCherry* forespores, consistent with the ∼3-fold reduction in heat-resistant spores in Δ*sipL*/*mCherry-sipL* relative to WT (**Figure S5**). Taken together, these results indicate that a C-terminal fusion of mCherry to SipL disrupts SipL-mCherry’s encasement of the forespore specifically in the absence of SpoVQ.

### Recombinant SpoVQ directly binds both SpoIVA and SipL

The localization (**Figures 5-6**) and double mutant analyses (**Figure 4**) supported a genetic interaction between SpoVQ and both SipL and SpoIVA, while the co-immunoprecipitation analyses revealed a biochemical interaction between SpoVQ, SipL, and SpoIVA during *C. difficile* sporulation (**Figure 1**). To test whether SpoVQ directly binds SipL and/or SpoIVA, we assessed whether immunoprecipitating FLAG-tagged SpoVQ would pull-down untagged SipL and/or SpoIVA in *C. difficile* sporulating cell lysates. For these analyses, we complemented Δ*spoVQ* with a strain producing C-terminally FLAG-tagged SpoVQ and co-immunoprecipitated SpoVQ-FLAG_3_ in the presence of detergent because SpoVQ is a bitopic membrane protein. Immunoprecipitation of FLAG-tagged SpoVQ failed to pull-down SipL, although small amounts of SpoIVA co-immunoprecipitated with SpoVQ*-*FLAG_3_, implying that SpoVQ may bind SpoIVA albeit weakly in the presence of NP-40 detergent (**Figure S6**).

Since our original co-immunoprecipitation analyses using FLAG-tagged SipL did not include detergent because SipL and SpoIVA are soluble proteins, we tested whether the inclusion of detergent would affect the efficiency of SpoVQ binding to SipL-FLAG_3_. While FLAG-tagged SipL pulled-down untagged SpoIVA (albeit at reduced levels) when NP-40 detergent was included, no SpoVQ was detected in the SipL-FLAG_3_ pull-down (**Figure S6**). In contrast, when detergent was not included in these analyses, trace amounts of SpoVQ were detected in the SipL-FLAG_3_ pull-downs (**Figure S6**). These findings suggested that the interaction between SipL and SpoVQ is either unstable in the presence of detergent or non-specific in the absence of detergent.

Despite these negative results, the genetic interactions we observed between SpoVQ, SpoIVA, and SipL (**Figure 4**) prompted us to use an alternative method to assess binding between SpoVQ and SipL and/or SpoIVA. Specifically, we used a recombinant co-affinity purification strategy that can detect binding between SpoIVA and SipL (20). This heterogeneous expression strategy allows proteins to be produced at higher levels and more readily measures direct binding. For these experiments, we co-produced His-tagged SpoVQ_Δ32_ (soluble domain) with either SipL or SpoIVA in *E. coli* and tested whether SipL or SpoIVA, respectively, would be present in the SpoVQ_Δ32_-His_6_ pull-downs. As a negative control, we used the soluble cysteine protease domain (CPD) derived from *Vibrio cholerae* MARTX toxin (46, 47). When His-tagged SpoVQ_Δ32_ was affinity-purified in the presence of either untagged SipL or SpoIVA, both proteins were enriched in the SpoVQ_Δ32_-His_6_ pull-downs. In contrast when His-tagged SpoVQ _Δ32_ wasaffinity-purified in the presence of untagged CPD, no enrichment of the CPD was observed.

These results indicate that recombinant SipL and SpoIVA can both interact with the soluble domain of SpoVQ at least in *E. coli* lysates.

## Discussion

Coat formation in *Clostridioides difficile* exhibits notable differences relative to *Bacillus subtilis* (10). For example, SipL is a clostridial-specific coat protein essential for coat localization to the forespore (20). Recent work has also identified Peptostreptococcaceae family-specific spore morphogenetic proteins that regulate *C. difficile* coat assembly in addition to other processes. CotL impacts *C. difficile* coat and cortex assembly (26), and CdeC is required for proper coat and exosporium assembly (48). These findings strongly suggest that the Peptostreptococcaceae family uses distinct pathways to assemble infectious spores (10).

In this study, we have identified SpoVQ as yet another Peptostreptococcaceae family-specific spore morphogenetic protein. Our analyses indicate that SpoVQ (CD3457) is a mother cell-specific bitopic membrane protein that specifically localizes to the outer forespore membrane (**Figure 5**) and regulates cortex thickness. The thinner cortex of regulation likely promotes higher levels of spontaneous germination (**Figure 3**). SpoVQ is also essential for cortex synthesis in Δ*spoIVA* and *sipL* mutants (**Figure 4**), which already exhibit abnormalities in cortex thickness and electron density (**Figure 4**, (23, 25)). Since SpoVQ directly binds both SpoIVA and SipL (**Figure 7**) and also influences SipL-mCherry localization to the forespore, (**Figure 6**) our data collectively suggest that SpoVQ functions to link coat and cortex assembly in *C. difficile*.

**Figure 7.**
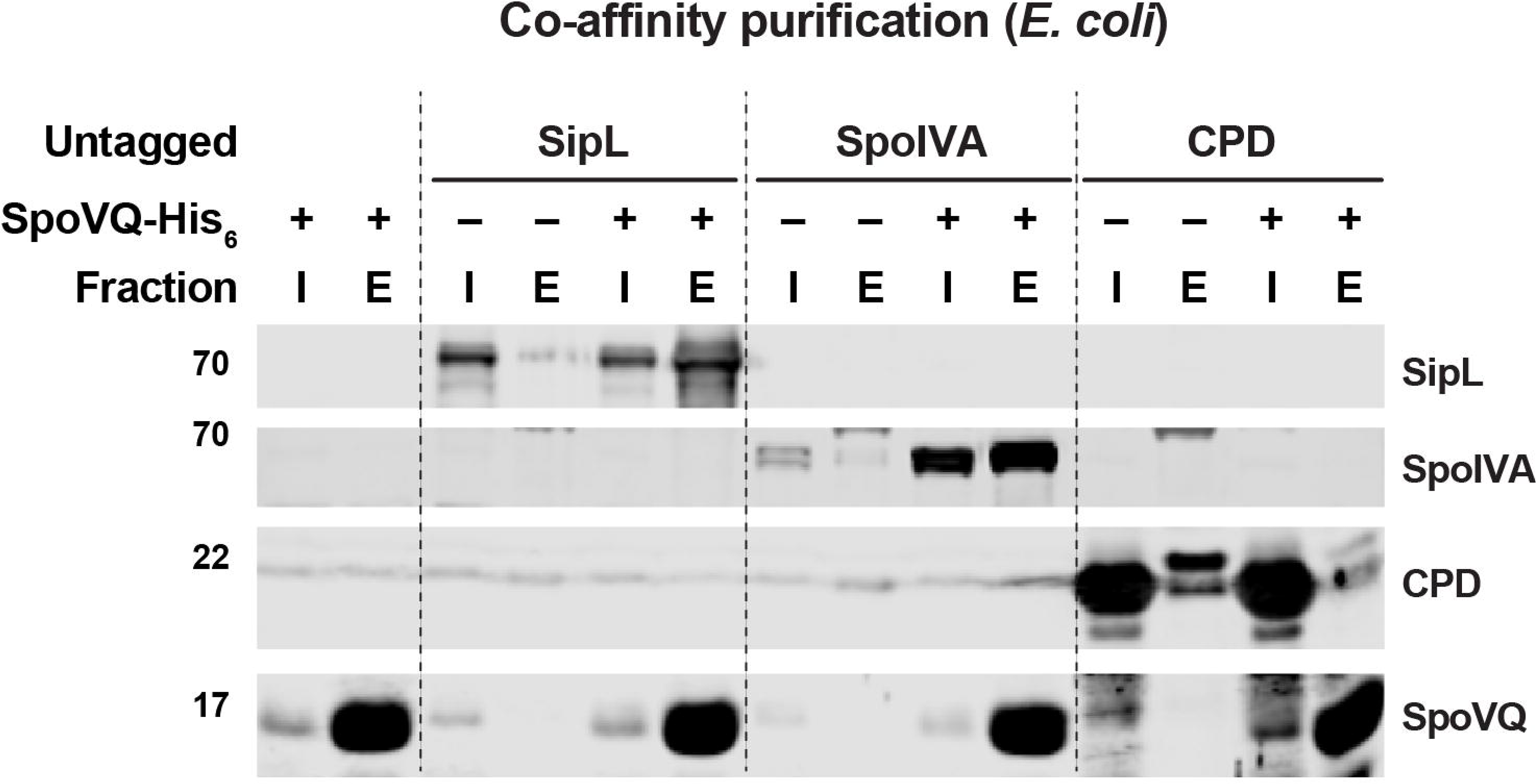
SpoVQ can separately bind SpoIVA and SipL in co-affinity purification analyses. Co-affinity purification analyses of His-tagged SpoVQ _Δ32_, which lacks its N-terminal transmembrane domain, with SipL, SpoIVA, and the CPD, a cysteine protease domain from *Vibrio cholerae* (46). The indicated proteins were co-produced in *E. coli*, and cleared soluble lysates were prepared. SpoVQ _Δ32_-His_6_ was affinity-purified using Ni-NTA resin, and high imidazole was used to elute SpoVQ _Δ32_-His_6_ and any associated proteins. Cleared lysate (input, I) and elution (E) samples were analyzed by Western blotting using anti-SipL, anti-SpoIVA, anti-CPD, and anti-SpoVQ antibodies.

A functional link between coat and cortex formation has long been established in *B. subtilis*, since *spoIVA* and *spoVM* mutants fail to make cortex and exhibit coat localization defects (16, 17, 21, 49). Recent work has shed light on the mechanism linking these two critical processes with the identification of SsdC as a Bacilli-specific, mother cell-derived membrane protein. SsdC localizes to the outer forespore membrane, regulates cortex synthesis, and genetically interacts with several basement layer coat morphogenetic proteins (50). While *B. subtilis* SsdC and *C. difficile* SpoVQ do not share sequence homology, they may share similar functions in linking cortex and coat formation through direct protein-protein interactions.

Addressing how SpoVQ impacts cortex formation in *C. difficile* is challenging given that the mechanisms controlling cortex synthesis in *C. difficile* remain poorly defined. Three of the enzymes critical for cortex peptidoglycan synthesis in *B. subtilis* are conserved in *C. difficile*, namely the SEDS glycosyltransferase-Class B penicillin-binding protein transpeptidase pair, SpoVE and SpoVD, and the SpoVB flippase (11, 12, 15, 51), but SpoVD is the only *C. difficile* protein that has been shown to regulate cortex synthesis to date (52). Analyzing the localization dependence of SpoVD and the remaining enzymes on *C. difficile* SpoVQ would address whether SpoVQ alters cortex thickness by impacting the localization and/or activity of these PG synthesizing enzymes. Studies directed at identifying binding partners of SpoVQ (beyond SpoIVA and SipL) using our functional epitope-tagged construct (**Figure S6**) may also provide insight into how SpoVQ modulates cortex synthesis.

Ironically, while we originally identified SpoVQ as a binding partner of SipL via co-immunoprecipitation (**Figure 1**), this interaction was not stable in the presence of detergent. Validating this interaction in the presence of detergent was necessary to rule out the possibility that SpoVQ indirectly binds SipL in the context of membrane micelles containing direct SipL interactors. Given that the stability of SpoIVA and SipL binding decreased markedly in the presence of detergent (**Figure S6**), it is possible that SipL and SpoVQ specifically bind during *C. difficile* spore formation but that the interaction is destabilized by detergent (**Figure 1** and **S6**). Future work should more rigorously assess direct binding between these proteins during *C. difficile* sporulation potentially by using cross-linking followed by co-immunoprecipitation analyses (53) or using co-localization analyses (54). Identifying the regions within recombinant SpoIVA and SipL that bind SpoVQ’s soluble domain and whether SpoVQ binds the SpoIVA-SipL complex more efficiently would also provide important insight into this interaction.

Another important question is whether SpoVQ’s soluble domain is oriented towards the intermembrane space or within the mother cell cytosol. TMHMM analyses predict relatively similar probabilities that SpoVQ’s C-terminal soluble domain is secreted vs. remains cytosolic (**Figure S1A**). Given the impact that SpoVQ has on cortex synthesis (**Figure 3**), it seems more likely that the C-terminal domain is secreted into the intermembrane space where it could regulate cortex synthesis. If this is the case, it raises the question of how SpoVQ, SpoIVA, and SipL physically interact, since SpoIVA and SipL are made in the mother cell cytosol. Recent work in *B. subtilis* suggests that some coat proteins can bridge the outer forespore membrane and bind cortex peptidoglycan. The *B. subtilis* coat morphogenetic protein, SafA, binds peptidoglycan (45) and the coat morphogenetic protein, SpoVID (55). Thus, SafA has a mechanism to reach across the outer forespore membrane despite lacking a predicted transmembrane domain. SafA binding to peptidoglycan depends on its N-terminal LysM domain (45), which is also required for SafA to bind SpoVID (55). Like *B. subtilis* SafA, *C. difficile* SipL carries a LysM domain that directly binds SpoIVA (20, 25), but whether SipL’s LysM domain binds peptidoglycan remains to be tested. Interestingly, SipL’s LysM domain is C-terminal, and loss of SpoVQ reduces the function of C-terminal mCherry fusion to SipL (**Figure 6**). Given that a C-terminal, but not N-terminal, fusion of mCherry to SipL disrupted SipL-mCherry localization in the absence of SpoVQ (**Figure 6**), SpoVQ would appear to require access to SipL’s C-terminal LysM domain. Thus, if SipL’s LysM domain binds peptidoglycan, SipL could provide a physical link between coat and cortex formation similar to how SafA “staples” the outer forespore membrane to the cortex (45). Addressing the accessibility of SpoVQ and SipL’s C-terminal domains to protease treatment in sporulating cells that have not completed engulfment would provide critical insight into these questions. These types of analyses will likely be challenging due to *C. difficile*’s S-layer reducing access to proteases and lysozyme (56) as well as the intrinsic resistance of its cell wall to lysozyme digestion (57).

Finally, another key question raised by our study is how SpoVQ specifically localizes to the outer forespore membrane. *spoVQ* expression is controlled by the mother cell-specific sigma factor, σ^E^ (30, 31), so SpoVQ could localize to all mother cell-derived membranes. However, we found that SpoVQ-mCherry localized to the polar septum during asymmetric division (**Figure 5**), so preferential localization to forespore membranes appears to occur early during spore formation (**Figure 5**). This localization is independent of whether the basement layer coat proteins, SpoIVA or SipL, are present, although slightly more cytosolic signal is observed in the absence of SpoIVA (**Figure 5**). These results contrast with the localization requirements for *B. subtilis* SsdC, whose preferential localization to the outer forespore membrane depends on basement layer coat proteins (50). SsdC also exhibits a different localization pattern than SpoVQ, since SsdC forms two distinct foci at the mother cell-forespore interface. Analyzing the localization dependence of SpoVQ-mCherry in mutants defective in engulfment, SpoIIQ-SpoIIIAH channel components, or cortex synthesis would provide insight into how SpoVQ specifically concentrates in the outer forespore membrane.

Taken together, considerable work remains to understand how the Peptostreptococcaceae family-specific spore morphogenetic protein SpoVQ regulates *C. difficile* cortex synthesis in concert with the coat morphogenetic proteins, SpoIVA and SipL. Regardless, our identification of this unique regulator reveals for the first time a direct link between coat localization and cortex synthesis in *C. difficile.* This work, along with recent work in *B. subtilis* (45, 50) suggests that the direct coordination of these two processes may be universally conserved across spore-forming bacteria even if the specific regulators and their mechanism of action may differ.

## Materials and Methods

### Bacterial strains and growth conditions

The *C. difficile* strains used are listed in Table S1 (Supplementary materials). All strains derive from the erythromycin-sensitive *630 erm pyrE* parental strain, which is a sequenced clinical isolate 630, which we used for *pyrE-*based allele coupled exchange (ACE) (29). Strains were grown on BHIS, brain heart infusion supplemented with yeast extract and cysteine (58) and taurocholate (TA, 0.1%, [wt/vol]; 1.9 mM), cefoxitin (8 µg/mL) and kanamycin (50 µg/mL) as needed. For ACE, *C. difficile* defined medium (59) (CDDM) was supplemented with 5-fluorootic acid (5-FOA at [2 mg/mL]) and uracil (at [5 µg/mL]).

The *Escherichia coli* strains used for HB101/pRK24-based conjugations and BL21(DE3) expression analyses are listed in Table S1. *E. coli* strains were grown at 37°C, shaking at 225 rpm in Luria-Bertani (LB) broth. The medium was supplemented with ampicillin (50 µg/mL), chloramphenicol (20 µg/mL) or kanamycin (30 µg/mL) as needed.

### *E. coli* strain construction

All primers used for cloning are listed in Table S2. Details of *E. coli* strain construction are provided in the Supplementary Text S1. Plasmid constructs were confirmed by sequencing using Genewiz and transformed into either the HB101/pRK24 conjugation strain (used with *C. difficile*) or BL21(DE3) expression strain.

### *C. difficile* strain construction and complementation

Allele-couple exchange (ACE) (29) was used to construct all complementation strains ACE was also used to introduce the *sipL* deletion Δ*erm*Δ*pyrE spoIVA* ATPase mutants. Complementations were performed as previously described by conjugating HB101/ pRK24 Δ*pyrE-*based strains (60) using allele-coupled exchange.

### Plate-based sporulation assay

*C. difficile* strains were grown overnight from glycerol stocks on BHIS plates containing with 0.1%, wt/vol taurocholate. Colonies from these plates were inoculated into BHIS liquid media and then back-diluted 1:25 once they were in stationary phase. The cultures were grown until they reached an optical density (OD_600nm_) between 0.4 - 0.7, and 120 µL was used to inoculate 70:30 agar plates (20). Sporulation was induced for 20 - 24 hours, after which cells were analyzed by phase-contrast microscopy to assess sporulation levels, harvested for western blot analyses, and the proportion of heat-resistant spores was measured.

### Immunoprecipitation analyses

Immunoprecipitations were performed on lysates prepared from cultures induced to sporulate on 70:30 plates for 24 h. The samples were processed as previously described (25), although for some of the replicates anti-FLAG magnetic beads (Sigma Aldrich) were used to pull-down FLAG-tagged proteins and any associated proteins rather than the Dynabead Protein G (Invitrogen) pre-bound with anti-FLAG M2 antibody (SigmaAldrich) as in (25). Where indicated, NP40 detergent was added at 0.1% (vol/vol) to the FLAG-IP buffer (FIB) (50 mM Tris, pH 7.5, 150 mM NaCl, 0.02% NaN_3_, 1 x protease inhibitor (HALT cocktail, ThermoScientific) after the lysate was prepared following cell lysis via bead-beating (Fastprep Pro, MP Biomedicals). Immunoprecipitations were performed on 2-3 biological replicates.

For immunoprecipitation experiments coupled to quantitative mass spectrometry-based proteomics, duplicate of each Δ*sipL/sipL* and Δ*sipL/sipL-FLAG_3_* strains were prepared and simultaneously. They were induced to sporulate for 24 h and the samples were prepared as previously described (25) using magnetic Dynabead Protein G (Invitrogen) coupled to anti-FLAG M2 antibody (Sigma Aldrich). Immunoprecipitations were carried out for 2 h at room temperature and the beads were washed four times with FIB. Efficient immunoprecipitation was confirmed by western blotting prior to proceeding with mass spectrometry.

The beads from the immunoprecipitations were washed an additional four times with PBS. The four different lysates of each replicate were resuspended in 90 uL digestion buffer (2M urea, 50 mM Tris HCl), and 2 ug of sequencing grade trypsin were added and shaken for 1 hour at 700 rpm. The supernatant was removed and placed in a fresh tube. The beads were then washed twice with 50 uL digestion buffer and combined with the supernatant. The combined supernatants were reduced (2 uL of 500 mM DTT, 30 minutes, RT), alkylated (4 uL of 500 mM IAA, 45 minutes, dark) and a longer overnight digestion was performed: 2 ug (4 uL) trypsin, and shaken overnight. The samples were then quenched with 20 uL of 10% Formic Acid and desalted on 10 mg Oasis cartridges. Desalted peptides from each pull-down were separately labeled with a specific iTR mass tagging reagent (lot A7024, Sciex) prior to mixing and analysis by LC-MS/MS. The tag layout of the iTRAQ 4-plex was as follows: 114, Control Rep 1; 115 Control Rep 2; 116, SIPL Rep 1; 117, SIPL Rep 2. Peptides were dissolved in 50 uL of 100% ethanol and one kit of iTRAQ reagent was added to each respective vial according to the lot information. The reaction proceeded for one hour at room temperature. The resulting pool of labeled peptides was dried and separated into six fractions using high pH (pH 10) fractionation on an SDB STAGE (Stop and go extraction) tip with 4 punches (Empore reversed phase extraction disks from 3M, SDB-XC reversed phase material, product number 2240/2340). The six fractions were as follows: Fraction 0, 3% acetonitrile; Fraction 1, 5% acetonitrile; Fraction 2, 10% acetonitrile; Fraction 3, 15% acetonitrile; Fraction 4, 20% acetonitrile; Fraction 5, 30% acetonitrile; Fraction 6, 50% acetonitrile.

Reconstituted peptides were separated on an online nanoflow EASY-nLC 1000 UHPLC system (Thermo Fisher Scientific) and analyzed on a benchtop Orbitrap Q Exactive plus mass spectrometer (Thermo Fisher Scientific). The peptide samples were injected onto a capillary μm diameter, New Objective, PF360–75-10-N-5) packed in-house with 20 cm C18 silica material (1.9 μ Maisch GmbH) and heated to 50 °C in column heater sleeves (Phoenix-ST) to reduce backpressure during UHPLC separation. Injected peptides were separated at a flow rate of 200 nL/min with a linear 230 min gradient from 100% solvent A (3% acetonitrile, 0.1% formic acid) to 30% solvent B (90% acetonitrile, 0.1% formic acid), followed by a linear 9 min gradient from 30% solvent B to 60% solvent B and a 1 min ramp to 90%B. Each sample fraction was run for 120 min, including sample loading and column equilibration times. The Q Exactive instrument was operated in the data-dependent mode acquiring HCD MS/MS scans (R=17,500) after each MS1 scan (R=70,000) on the 12 top most abundant ions using an MS1 ion target of 3× 10^6^ ions and an MS2 target of 5×10^4^ ions. The maximum ion time utilized for the MS/MS scans was 120 ms; the HCD-normalized collision energy was set to 27; the dynamic exclusion time was set to 20s, and the peptide match and isotope exclusion functions were enabled.

All mass spectra were processed using the Spectrum Mill software package v6.0 pre-release (Agilent Technologies) which includes modules developed by us for iTR -based quantification. For peptide identification MS/MS spectra were searched against human Uniprot database to which a set of common laboratory contaminant proteins was appended. Search parameters included: ESI-QEXACTIVE-HCD scoring parameters, trypsin enzyme specificity with a maximum of two missed cleavages, 40% minimum matched peak intensity, +/− 20 ppm precursor mass tolerance, +/− 20 ppm product mass tolerance, and carbamidomethylation of cysteines and iTR labeling of lysines and peptide n-termini as fixed modifications. Allowed variable modifications were oxidation of methionine, N-terminal acetylation, Pyroglutamic acid (N-termQ),Deamidated (N),Pyro Carbamidomethyl Cys (N-termC),with a precursor MH+ shift range of −18 to 64 Da. Identities interpreted for individual spectra were automatically designated as valid by optimizing score and delta rank1-rank2 score thresholds separately for each precursor charge state in each LC-MS/MS while allowing a maximum target-decoy-based false-discovery rate (FDR) of 1.0% at the spectrum level.

The expectation maximization algorithm was applied to the results of the peptide report. The list of most likely observed proteins was generated for each channel of the MS experiment based on Swiss-Prot and TrEMBLE databases of protein sequences. Next, ratios of intensities between channels were calculated and median normalized. Resulting data were analyzed and visualized using R. Statistical analyses were performed via moderated t-test from R package limma to estimate p values for each protein and the false discovery rate corrections (FDR) were applied to account for multiple hypothesis testing. Plots were created using in-house written R scripts and gplot2. Hits were defined as proteins with a ≥ 2-fold enrichment with a p-value < 0.01.

### Heat resistance assay on sporulating cells

Heat-resistant spore formation was assessed 20-24 hrs after sporulation induction as previously described (33). Heat-resistance efficiencies represent the average ratio of heat-resistant colony forming units obtained from spores in a given sample relative to the average ratio determined for the wild type based on a minimum of three biological replicates. Statistical significance was determined using one-way ANOVA and Tukey’s test.

### Western-blot analyses

Samples for Western-blotting analyses were prepared as previously described (20). Briefly, sporulating cell pellets were resuspended in 100 μL of PBS and subjected to three freeze-thaw cycles. Denaturing EBB buffer was added to the sample at ∼1:1 ratio (8 M urea, 2 M thiourea, 4% (wt/vol) SDS, 2% (vol/vol) β were boiled for 20 min, vortexed, pelleted at high-speed, and resuspended in the same supernatant to maximize protein solubilization. Finally, the samples were boiled for 5 minutes and pelleted again at high-speed. Samples were resolved on SDS-PAGE gels, transferred to Immobilon-FL polyvinylidene difluoride (PVDF) membranes, and blocked in Odyssey Blocking Buffer with 0.1% (vol/vol) Tween 20. Rabbit anti-SpoVQ _Δ32_ (this study) and anti-CPD (46) antibodies were used at 1:1,000 dilution. Rabbit anti-SipL _LysM_ and mouse anti-SpoIVA (61) antibodies were used at a 1:2,500 dilution. Rabbit anti-mCherry (Abcam, Inc.) was used at a 1:2,000 dilution. IRDye 680CW and 800CW infrared dye-conjugated secondary antibodies were used at a 1:20,000 dilution, and blots were imaged on an Odyssey LiCor CLx imaging system.

### Spore purification

Sporulation was induced in 70:30 agar plates for 2 to 3 days as previously described (18, 62). *C. difficile* sporulating cultures were lysed in 5-6 cycles of ice-cold water washes, incubated on ice overnight at 4°C, pelleted and treated with DNAse I (New England Biolabs) at 37°C for 60 minutes. Finally, spores were purified on a 20%:50% HistoDenz gradient (Sigma-Aldrich) and resuspended in water. Spore purity was determined by phase-contrast microscopy (>95%), and the optical density of the spore preparation was measured at OD_600_. Spore yields were quantified by measuring the OD_600nm_ of the spore purifications from four 70:30 plates per replicate. The average of six biological replicates was calculated and statistical significance was determined using a one-way ANOVA and Tukey’s test. Spores were stored in water at 4°C.

### Germination assays

Germination assays were performed as previously described (63). Approximately, ∼1 x 10^7^ spores (0.35 OD_600_ units) were resuspended in 100 µl of water, and 10 µL of this mixture was removed for 10-fold serial dilutions in PBS. The dilutions were plated on either BHIS alone or BHIS containing 0.1% taurocholate germinant. Colonies arising from germinated spores were enumerated after 18-24 hrs. Germination efficiencies were calculated by averaging the CFUs produced by spores for a given strain relative to the number produced by wild-type spores from at least three independent spore preparations. Statistical significance was determined by performing a one-way analysis of variance (ANOVA) on natural log-transformed data using Tukey’s test. The data were transformed because the use of independent spore preparations resulted in a non-normal distribution for spontaneous germination results.

### TEM analysis

Sporulating cultures (23-24 h) were fixed and processed for electron microscopy by the University of Vermont Microscopy Center according to previously published protocols (20). A minimum of 50 full-length sporulating cells were used for phenotype counting.

### mCherry fluorescence microscopy

Live-cell fluorescence microscopy was performed using Hoechst 33342 (15 µg/mL; Molecular Probes) and mCherry protein fusions. Samples were prepared on agarose pads using Gene Frames (ThermoScientific) as previously described (23). Images were taken 30 minutes after harvesting the *C. difficile* sporulating cultures to allow the mCherry fluorescent signal to reconstitute under aerobic conditions.

Phase-contrast and fluorescence microscopy were carried out on a Nikon 60x oil immersion objective (1.4 numerical aperture [NA]) using a Nikon 90i epifluorescence microscope. A CoolSnap HQ camera (Photometrics) was used to acquire multiple fields for each sample in 12-bit format with 2-by-2 binning, using NIS-Elements software (Nikon). The Texas Red channel was used to acquire images after 300 ms for mCherry-SpoIVA; 100 ms for SipL-mCherry; and 400 ms for CotE-mCherry. Hoechst stain was visualized using 90-ms exposure time and 3-ms exposure was used for phase-contrast pictures. Finally, 10 MHz images were imported to Adobe Photoshop CC 2019 software for pseudocoloring and minimal adjustments in brightness and contrast levels. Protein localization analysis were performed on a minimum of three independent biological replicates.

### Protein purification for antibody production and co-affinity purification analyses

His_6_-tagged proteins were affinity-purified from *E. coli* BL21(DE3) cultures (see **Table S2** in Supplementary Materials) grown in 2YT media (5 g NaCl, 10 g yeast extract, and 15 g tryptone per liter) after an overnight induction with 250 µM IPTG at 19°C as previously described (62). The anti-SpoVQ antibody was raised against SpoVQ _Δ32_-His_6_, which lacks its transmembrane domain (first 32 codons), and the antibody against CotL was raised against CotL-His_6_ in rabbits by Cocalico Biologicals (Reamstown, PA).

Cultures were pelleted, resuspended in lysis buffer (500 mM NaCl, 50 mM Tris [pH 7.5], 15 mM imidazole, 10% [vol/vol] glycerol), flash frozen in liquid nitrogen, and sonicated. The insoluble material was pelleted, and the soluble fraction was incubated with Ni-NTA agarose beads (0.5 ml; Qiagen) for 3 hrs, and eluted using high-imidazole buffer (500 mM NaCl, 50 mM Tris [pH 7.5], 200 mM imidazole, 10% [vol/vol] glycerol) after nutating the sample for 5-10 min. The resulting induction and eluate fractions were separated by SDS-PAGE and analyzed by Western blotting as described above.

For the co-affinity purification assays, *E. coli* BL21(DE3) strains were simultaneously transformed with two plasmids to express combinations of SpoVQ _Δ32_-His_6_ and either untagged SpoIVA, SipL, or CPD, or empty vector. Co-affinity purifications were performed using Ni^2+^-affinity resin as described above.

## Acknowledgments

We would like to thank N. Bishop, M. Norris, and J. Schwarz for excellent assistance in preparing samples for transmission electron microscopy throughout this study; A. Camilli for access to the Nikon microscope; N. Minton (U. Nottingham) for generously providing us with access to the 630 Δ*erm*Δ*pyrE* strain and pMTL-YN1C and pMTL-YN3 plasmids for allele-coupled exchange (ACE). Research in this manuscript was funded by R01AI22232 from the National Institutes of Allergy and Infectious Disease (NIAID) to A.S. A.S. is a Burroughs Wellcome Investigator in the Pathogenesis of Infectious Disease supported by the Burroughs Wellcome Fund. The content is solely the responsibility of the author(s) and does not necessarily reflect the views of the Burroughs Wellcome, NIAID, or the National Institutes of Health. The funders had no role in study design, data collection and interpretation, or the decision to submit the work for publication.

