## Supplementary Materials for "Identification of a novel regulator of *Clostridioides difficile* cortex formation"

**Supplementary Material Description**

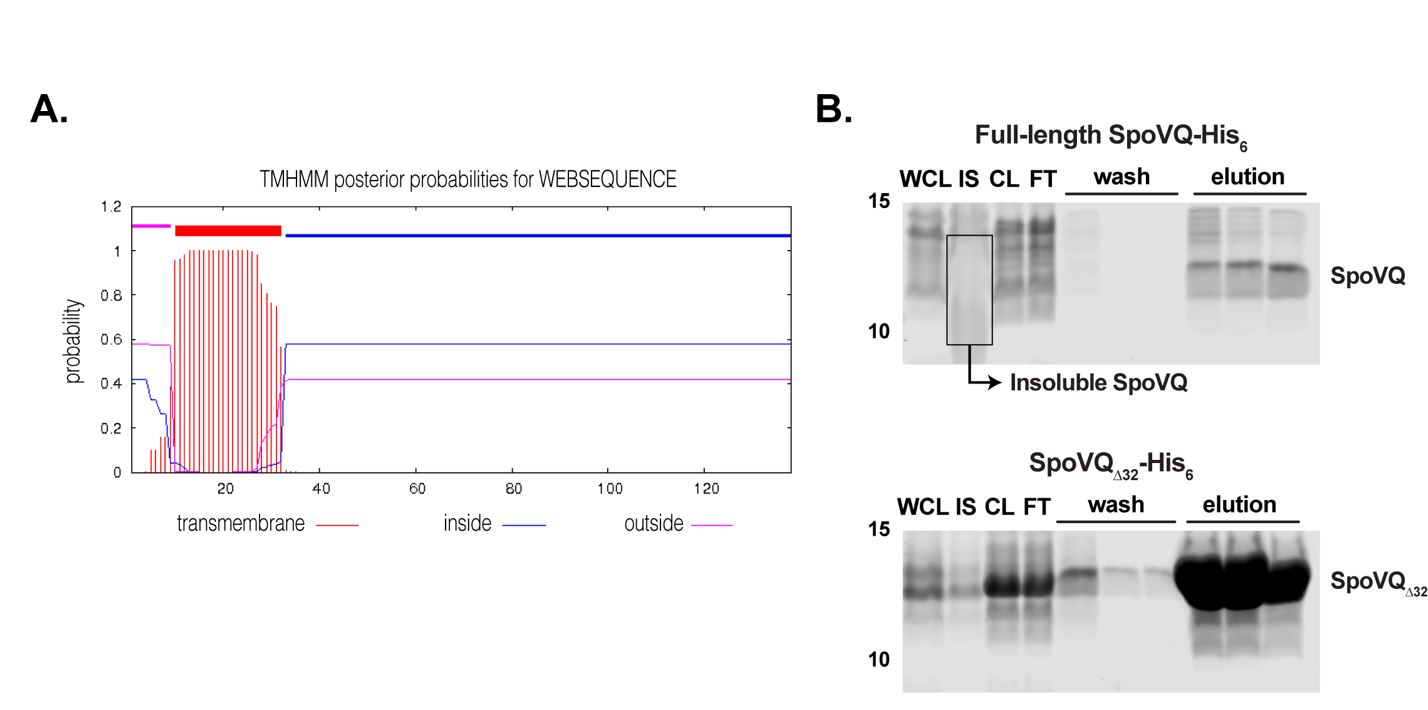
**Figure S1. SpoVQ has an N-terminal transmembrane domain.** (A) TMHMM prediction of transmembrane helices in SpoVQ. Topology prediction regarding whether the C-terminal soluble domain is secreted or cytosolic is ambiguous. (B) Ni^2+^-affinity purification of full-length His-tagged SpoVQ and His-tagged SpoVQ_∆32_, which is missing its N-terminal 32 aa (predicted transmembrane domain). Whole cell lysate (WCL), insoluble (IS), cleared lysate (soluble), flow-through (FT), wash, and imidazole elution fractions from the purification were resolved by SDS-PAGE and Coomassie stained. Deletion of the transmembrane domain markedly increases the solubility of SpoVQ.

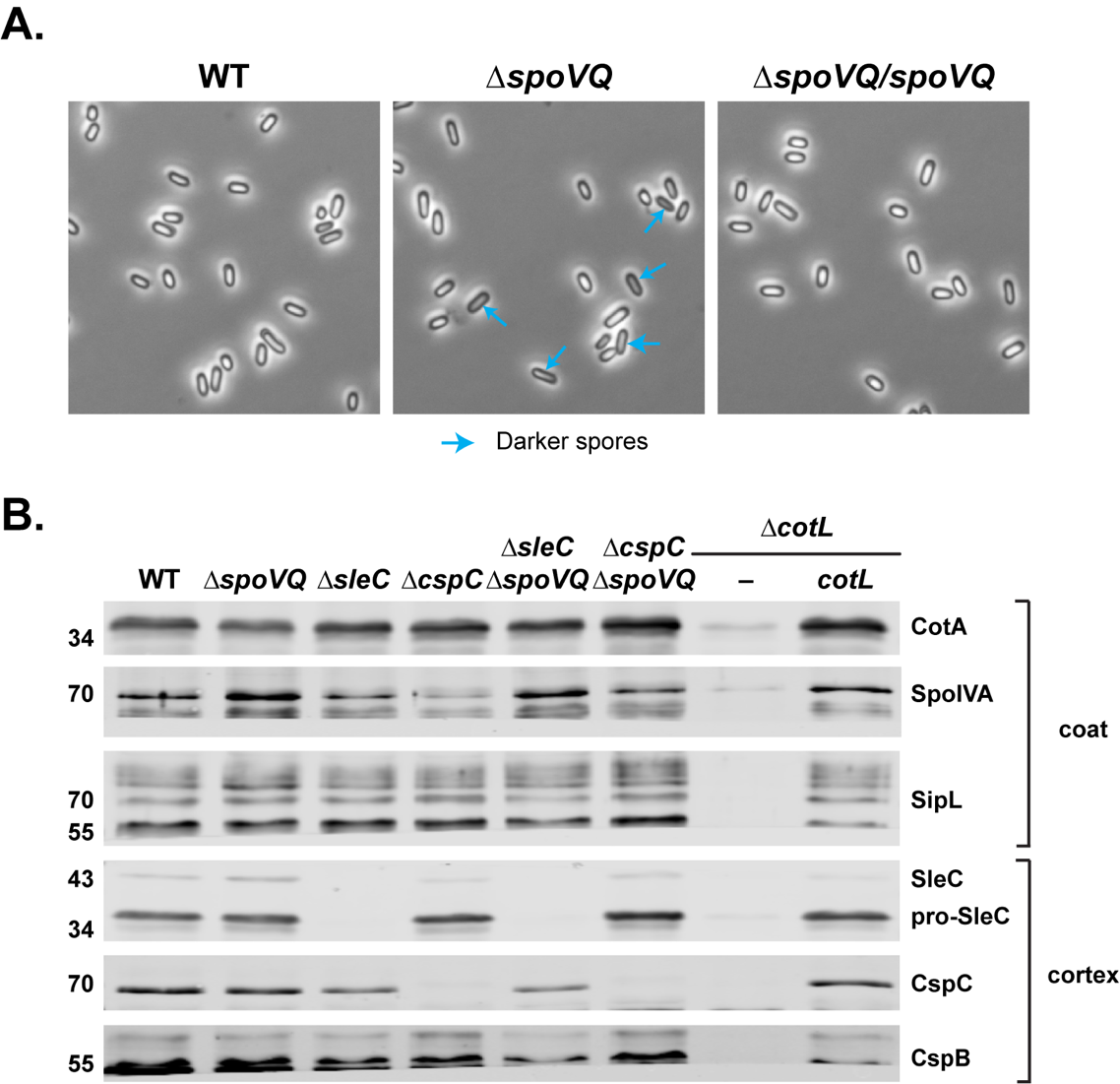

**Figure S2. ∆*spoVQ* spore characterization.** (A) Phase-contrast microscopy of purified wild-type (WT), ∆*spoVQ*, and ∆*spoVQ* complementation spores. Blue arrows highlight spores that are slightly darker than other spores. (B) Western blot analyses of ∆*spoVQ,* ∆*cotL*, and germination mutant spores. CotA, SpoIVA, and SipL are coat proteins (1, 2), while SleC , CspC, and CspB are all predicted to be cortex-localized (3, 4). *C. perfringens* SleC has been localized to the cortex region using immunogold labeling previously (5). Western blots are representative of three biological replicates.

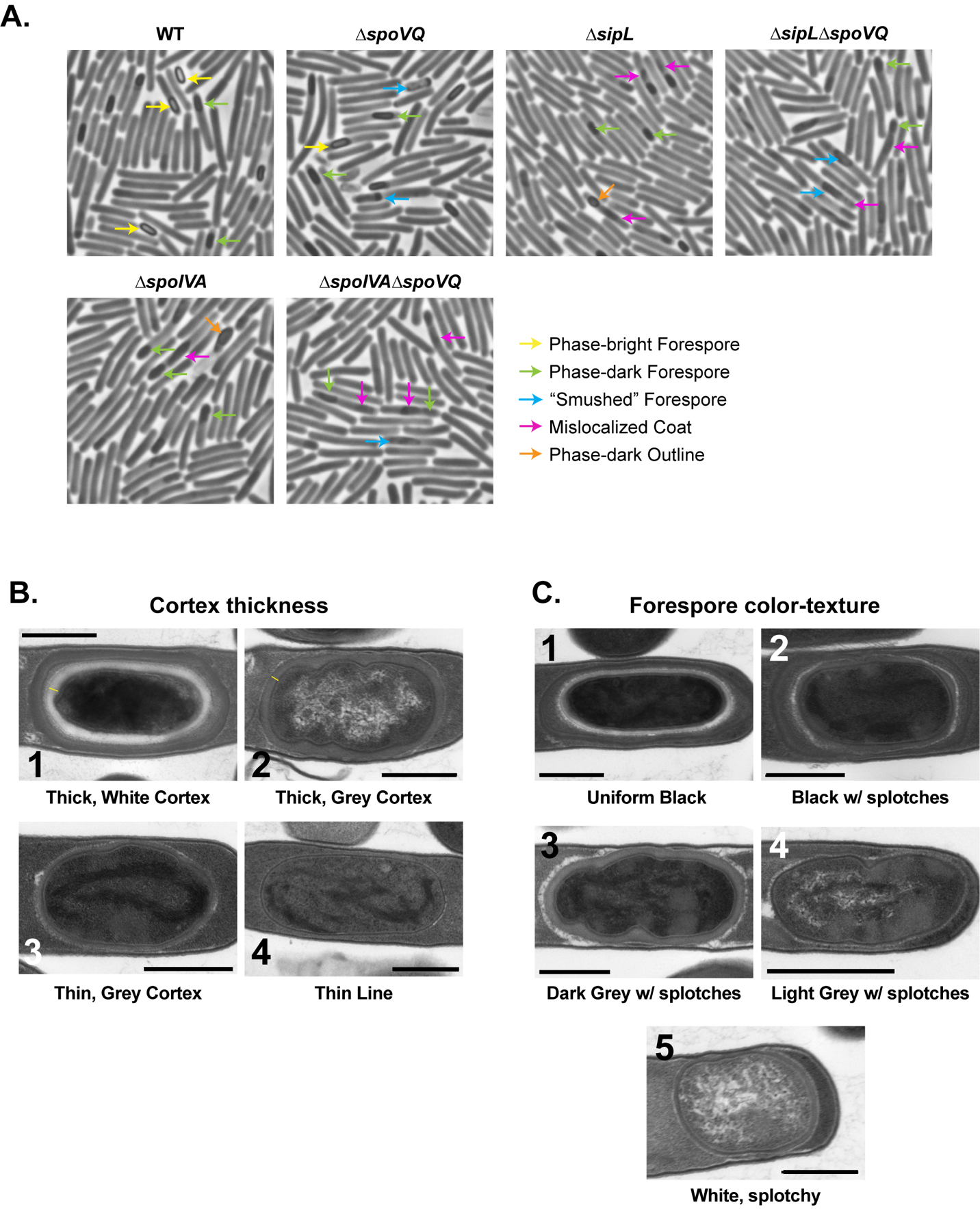
**Figure S3. Morphological analyses of sporulating cultures of ∆*spoVQ*, ∆*sipL*, and ∆*spoIVA* single and double mutants**. (A) Phase-contrast microscopy analyses. Phase-bright forespores (yellow arrows) are detected in wild-type and ∆*spoVQ* spores but not in ∆*sipL* or ∆*spoIVA* strain backgrounds. “Smushed” forespores (blue arrows) are visible in strains lacking *spoVQ*, while mislocalized coat (pink arrows) is visible in strains lacking either *sipL* or *spoIVA*. ∆*sipL* and ∆*spoIVA* strains occasionally make forespores with a distinct phase-dark outline (orange arrows) that likely represents synthesized cortex. (B and C) Transmission electron microscopy analyses of sporulating culture shown in (A). Examples of the phenotypes scored for cortex thickness (B) and forespore color/texture (C) are shown. Thin yellow lines in (B) highlight the cortex region where detectable. Scale bars represent 500 nm.

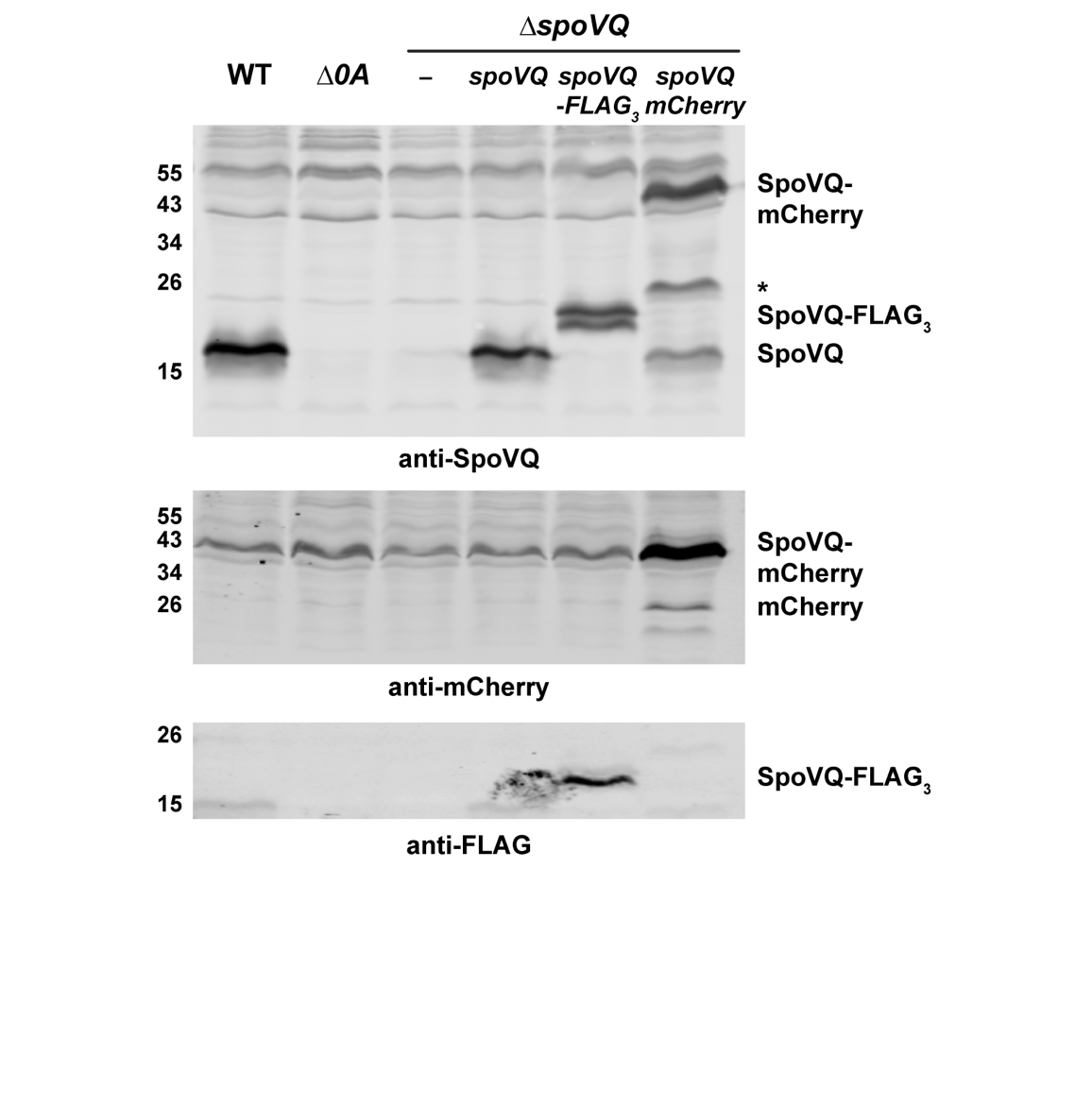

**Figure S4. Western blot analyses of tagged *spoVQ* strains**. *spoVQ-FLAG_3_* and *spoVQ-mCherry* encode C-terminal epitope and fluorescent protein fusions to SpoVQ, respectively. Asterisk represents a cleavage product of SpoVQ-mCherry. Some free mCherry is detected. The antibodies used for each blot are shown below. The result is representative of at least two biological replicates.

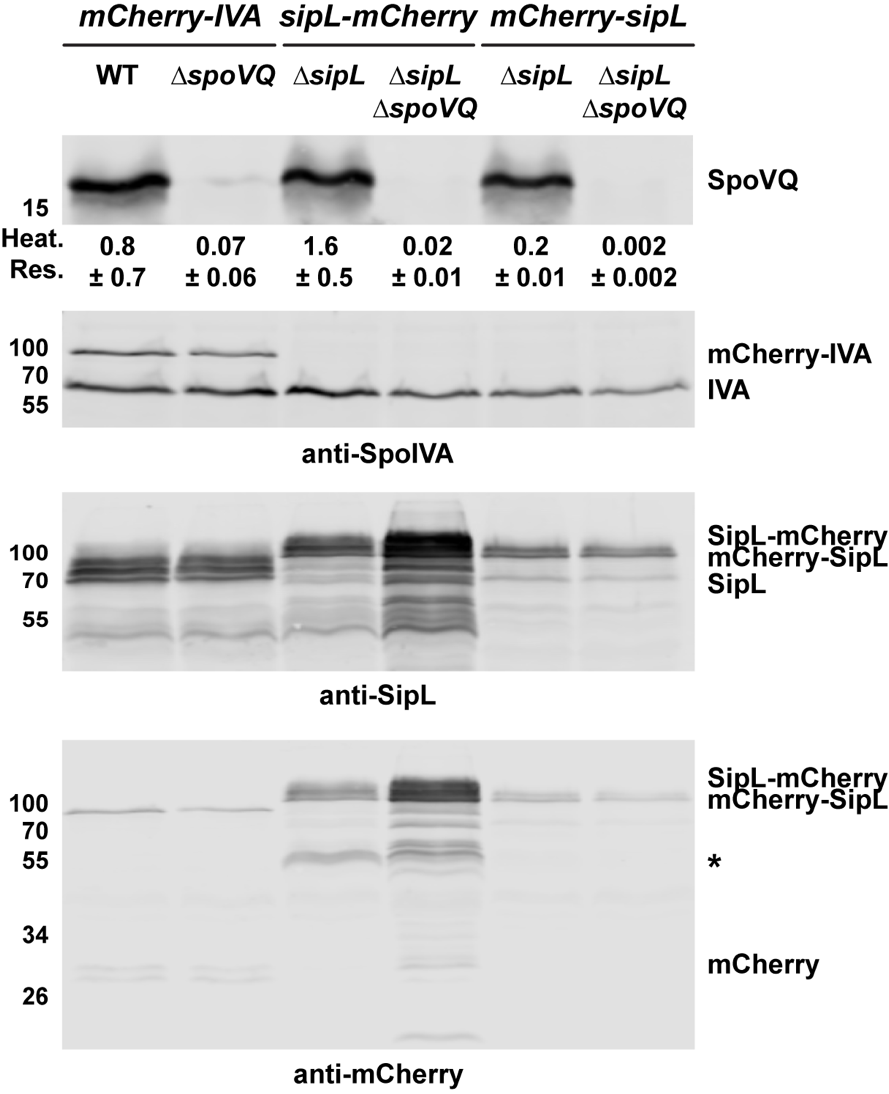

**Figure S5. Western blot analyses of strains carrying mCherry-fusions to SipL and SpoIVA**. *mCherry-spoIVA (IVA*) was conjugated into a wild-type and ∆*spoVQ* strain background to create merodiploid strains. *sipL-mCherry* (6) and *mCherry-sipL* constructs were conjugated into a ∆*sipL* strain so that the fusion protein is the only SipL version present. Functionality of the fusion proteins was assessed using a heat resistance assay (Heat res., (7)). The results are based on a minimum of three biological replicates (with the exception of ∆*sipL∆spoVQ*/*mCherry-sipL*, which is based on two biological replicates). (∆*sipL/sipL-mCherry* vs. ∆*sipL∆spoVQ/sipL-mCherry*, p < 0.005; ∆*sipL/sipL-mCherry* vs. ∆*sipL∆spoVQ/ mCherry-sipL*, p < 0.005). Minimal degradation of fluorescent protein fusions was observed. It should also be noted that SipL-mCherry and mCherry-SipL are virtually identical in size. Results are representative of two biological replicates.

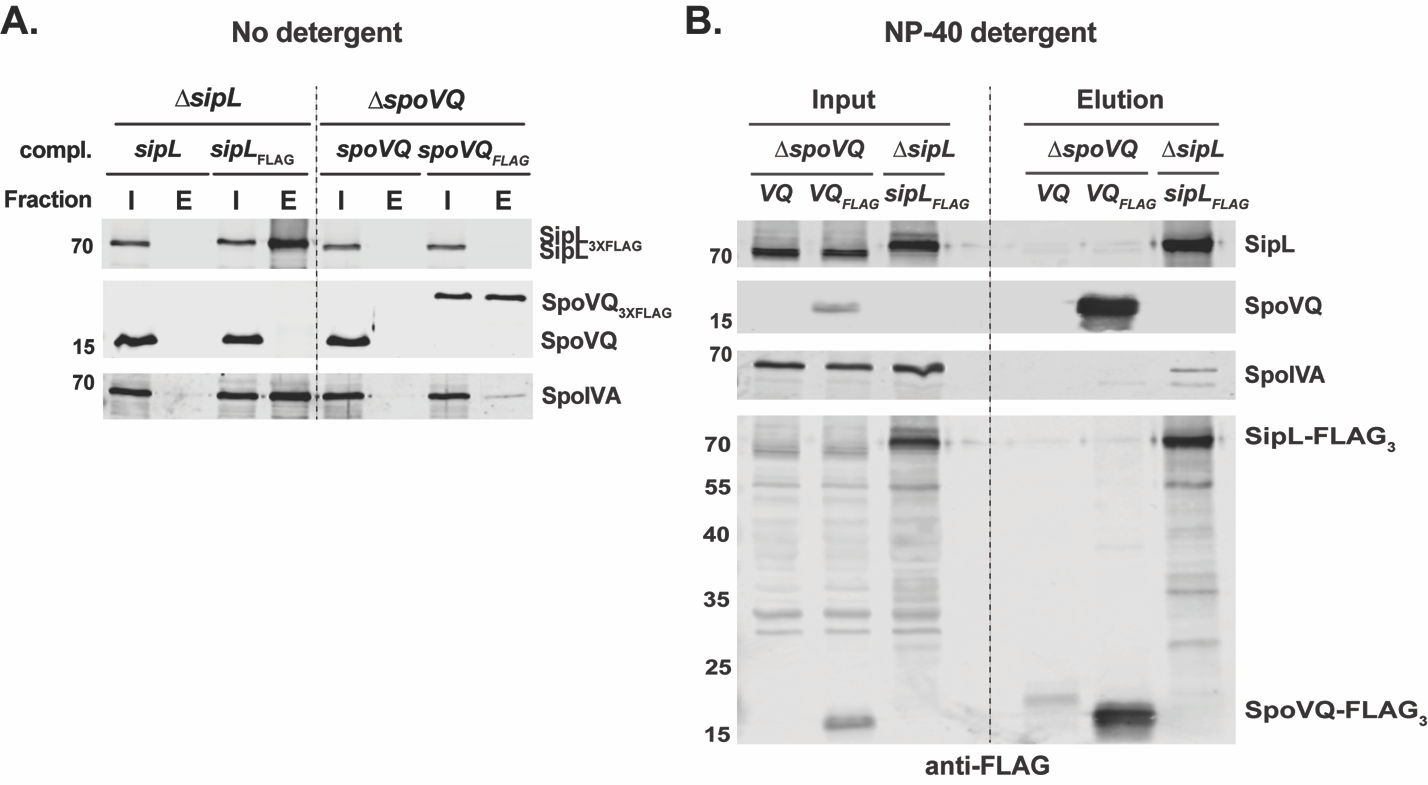

**Figure S6**. **Co-immunoprecipitation analyses of FLAG-tagged *sipL* and *spoVQ* from *C. difficile* sporulating cell lysates.** (A) FLAG-tagged SipL and SpoVQ were immunoprecipitated from cleared lysates prepared from the indicated *C. difficile* ∆*sipL* and ∆*spoVQ* complementation strains, respectively, using anti-FLAG magnetic beads. No detergent was present in these analyses. (I = Input fraction) After several washes, proteins retained by the beads were eluted using FLAG peptide (E = Elution fraction). The untagged *sipL* and *spoVQ* complementation strains serve as negative controls to assess specific binding of FLAG-tagged proteins to untagged beads. Sporulation was induced for 24 hrs before lysates were prepared. Untagged SpoIVA was pulled-down robustly by SipL-FLAG_3_, while untagged SpoVQ co-purified with SipL-FLAG_3_ with lower apparent efficiency. The immunoprecipitations shown are representative of three independent biological replicates. (B) SipL-FLAG_3_ and SpoVQ-FLAG_3_ co-immunoprecipitations were performed in the presence of 0.1% NP40 detergent and analyzed similarly to (A). Lower levels of untagged SpoIVA were co-purified with SipL-FLAG_3_ and SpoVQ-FLAG_3_, respectively.

**Supplementary Text S1 – *E. coli* strain construction**

**pET28a-*spoVQ*-His_6_**. To clone full-length His-tagged SpoVQ for protein expression in *E. coli*, primer pair #2642 and 2643 were used to amplify the *spoVQ* gene lacking its stop codon. The resulting PCR product was digested with NcoI and XhoI and ligated into pET28a digested with the same enzymes. The ligation was transformed into DH5α, and the resulting construct was sequenced verified before transforming the plasmid into BL21(DE3) for protein expression.

**pET28a-*spoVQ_∆_*_32_-His_6_**. To clone His-tagged SpoVQ lacking its N-terminal transmembrane domain (codons 1-32) for protein expression in *E. coli*, primer pair #2464 and 2465were used to amplify the *spoVQ* gene lacking its stop codon and first 32 codons. The resulting PCR product was digested with NcoI and XhoI and ligated into pET28a digested with the same enzymes. The ligation was transformed into DH5α, and the resulting construct was sequenced verified before transforming the plasmid into BL21(DE3) for protein expression.

**pET22b-*cotL*-His_6_**. To clone full-length His-tagged CotL for protein expression in *E. coli*, primer pair #2946 and 2947 were used to amplify the *cotL* gene lacking its stop codon. The resulting PCR product was digested with Nde and XhoI and assembled into pET22b digested with the same enzymes using Gibson assembly. The assembly was transformed into DH5α, and the resulting construct was sequenced verified before transforming the plasmid into BL21(DE3) for protein expression.

**pET22b-CPD(TAA)**. To clone a His-tagged *Vibrio cholerae* cysteine protease domain (CPD) from the MARTX toxin gene, primer pair #2573 and 2574 were used to amplify the region encoding the CPD (aa 3439-3650) and to add a stop codon. The resulting PCR product was digested with NdeI and XhoI and Gibson assembled into pET22b digested with the same enzymes. The assembly was transformed into DH5α, and the resulting construct was sequenced verified before transforming the plasmid into BL21(DE3) for protein expression.

**pMTL-YN3 ∆*spoVQ***. Primer pair #2484 and 2486 were used to amplify the region 704 bp upstream of the *spoVQ* gene along with the first 22 codons off *C. difficile* genomic DNA. Primer pair #2485 and 2487 were used to amplify the region 665 bp downstream of the *spoVQ* gene and the last 12 codons of *spoVQ* using *C. difficile* genomic DNA as the template. The PCR products resulting PCR products were cloned into pMTL-YN3 digested with AscI and SbfI using Gibson assembly.

**pMTL-YN1C *spoVQ****.* To clone the *spoVQ* complementation construct, primer pair #2540 and 2543 was used to amplify the *spoVQ* gene including the stop codon and the region 232 bp upstream of *spoVQ*. The resulting PCR product was cloned into pMTL-YN1C digested with NotI and XhoI using Gibson assembly.

**pMTL-YN1C *spoVQ-FLAG_3_***. To clone the *spoVQ* complementation construct encoding a C-terminal FLAG_3_ epitope tag, primer pair #2540 and 2544 was used to amplify the *spoVQ* gene without the stop codon and its promoter region (232 bp upstream) along with sequence encoding part of a FLAG epitope. The resulting PCR product was assembled with the following g-block encoding the FLAG_3_ epitope into pMTL-YN1C along with digested with NotI and XhoI using Gibson assembly.

gBlock: Cdif CD3457-3xFLAG for pMTL-YN1C

GTATAGAAATGGACAAAATAGAAAGTGTGGTTGAGGAAGGAAAGGATGTTTTGAGAGTCAAGATAAAGTATAAAGATAAGGAAGATTCCTTTCCATACATAGTCGTTGAAACAAATATGAGTGAACTTCCAGATAGAATAGAATTAAATGCAACGAAAGATTATAAAGATGATGATGATAAAGACTATAAAGATGACGATGATAAGGATTATAAGGATGATGATGACAAATAACTCGAGGCCTGCAGACATGCAAGCTTGGCACTG

**pMTL-YN1C *spoVQ-mCherry***. To generate a construct encoding a C-terminal mCherry fusion to SpoVQ, primer pair #2540 and 2541 was used to amplify *spoVQ* and its promoter region (232 bp upstream) along with part of the *mCherry* gene*.* Primer pair #2542 and #2133 was used to amplify a codon-optimized *mCherry* gene as the template (8). The resulting PCR products were gel purified and assembled into pMTL-YN1C using Gibson assembly.

**pMTL-YN1C *spoVQ*_∆32_**. To generate a construct encoding a SpoVQ lacking its transmembrane domain spanning aa 1-32, primer pair #2540 and 2549 was used to amplify *spoVQ* and its promoter region (232 bp upstream) along with part of the *mCherry* gene*.* Primer pair #2550 and #2543 was used to amplify *spoVQ* (spanning codons 33 through its stop codon). The resulting PCR products were assembled into pMTL-YN1C using Gibson assembly.

**pMTL-YN3 ∆*cotL***. Primer pair #2820 and 2822 were used to amplify the region 763 bp upstream of the *cotL* gene along with the first 12 codons off *C. difficile* genomic DNA. Primer pair #2821 and 2823 were used to amplify the region 813 bp downstream of the *cotL* gene and the last 9 codons of *cotL* using *C. difficile* genomic DNA as the template. The PCR products resulting PCR products were cloned into pMTL-YN3 digested with AscI and SbfI using Gibson assembly.

**pMTL-YN1C *cotL****.* To clone the *cotL* complementation construct, primer pair #2833 and 2834 was used to amplify the *cotL* gene including the stop codon and the region 327 bp upstream of *cotL*. The resulting PCR product was cloned into pMTL-YN1C digested with NotI and XhoI using Gibson assembly.

**pMTL-YN1C *mCherry-sipL***. To generate a construct encoding an N-terminal mCherry fusion to SipL, primer pair #2165 and 3022 was used to amplify the promoter region of *sipL.* Primer pair #3021 and #3020 was used to amplify a codon-optimized *mCherry* gene as the template (8). Primer pair #3019 and 2166 was used to amplify the *sipL* gene including the stop codon off *C. difficile* genomic DNA. The resulting PCR products were gel purified and assembled into pMTL-YN1C using Gibson assembly.

**Supplementary Table S1. Strains used in this study**

| **Strain**  **#** | **Strain name** | **Relevant genotype or features** | | **Source/reference** | | |
| --- | --- | --- | --- | --- | --- | --- |
| ***C. difficile* strains – 630∆*erm*** | | | | |  | |
| 803 | 630∆*erm*∆*pyrE* ∆*spoIVA* | | 630∆*erm* ∆*pyrE* with *spoIVA* (*CD2629*) deleted | | | (9) |
| 846 | 630∆*erm*-p | | *erm*-sensitive derivate of 630 with *pyrE* restored | | | (10) |
| 849 | 630∆*erm ∆spo0A*-p | | 630∆*erm* ∆*spo0A* with *pyrE* restored | | | (10) |
| 880 | 630∆*erm ∆spoIVA*-p | | 630∆*erm* ∆*spoIVA* with *pyrE* restored | | | (9) |
| 925 | 630∆*erm ∆sleC*-p | | 630∆*erm* ∆*sleC* with *pyrE* restored | | | (10) |
| 1005 | 630∆*erm*∆*pyrE* ∆*sipL* | | 630∆*erm*∆*pyrE* with *sipL* deleted | | | (11) |
| 1010 | 630∆*erm* ∆*sipL*-p | | 630∆*erm* with *sipL* deleted and *pyrE* restored | | | (11) |
| 1013 | 630∆*erm* ∆*sipL*/*sipL* | | 630∆*erm* ∆*sipL* with *sipL* in the *pyrE* locus | | | (11) |
| 1144 | 630∆*erm*/*mCherry*-*IVA* | | 630∆*erm* with *mCherry-IVA* in the *pyrE* locus | | | (9) |
| 1158 | 630∆*erm ∆sipL*/*sipL-mCherry* | | 630∆*erm* ∆*sipL* with *sipL-mCherry* in the *pyrE* locus | | | (11) |
| 1238 | 630∆*erm ∆cspC-p* | | 630∆*erm* ∆*cspC* with *pyrE* restored | | | (10) |
| 1377 | 630∆*erm ∆sipL*/*sipL-FLAG_3_* | | 630∆*erm ∆sipL* with *sipL-FLAG_3_* in the *pyrE* locus | | | (6) |
| 1797 | 630∆*erm ∆spoVQ*∆*pyrE* | | 630∆*erm* ∆*pyrE* with *spoVQ* deleted | | | This study |
| 1804 | 630∆*erm ∆spoVQ-p* | | 630∆*erm* ∆*spoVQ* with *pyrE* restored | | | This study |
| 1807 | 630∆*erm ∆spoVQ/spoVQ* | | 630∆*erm* ∆*spoVQ* with *spoVQ* in the *pyrE* locus | | | This study |
| 1810 | 630∆*erm ∆spoVQ/spoVQ_-_FLAG_3_* | | 630∆*erm* ∆*spoVQ* with *spoVQ-FLAG_3_* in the *pyrE* locus | | | This study |
| 1813 | 630∆*erm* ∆*spoVQ/spoVQ-mCherry* | | 630∆*erm* ∆*spoVQ* with *spoVQ-mCherry* in the *pyrE* locus | | | This study |
| 1825 | 630∆*erm ∆spoVQ/spoVQ_∆32_* | | 630∆*erm* ∆*spoVQ* with *spoVQ_∆32_* in the *pyrE* locus | | | This study |
| 1888 | 630∆*erm ∆spoVQ*/*mCherry-IVA* | | 630∆*erm ∆spoVQ* with *mCherry-spoIVA* in the *pyrE* locus | | | This study |
| 1929 | 630∆*erm* ∆*cspC∆spoVQ*∆*pyrE* | | 630∆*erm* ∆*cspC∆pyrE* with *spoVQ* deleted | | | This study |
| 1932 | 630∆*erm* ∆*sleC∆spoVQ*∆*pyrE* | | 630∆*erm* ∆*sleC∆pyrE* with *spoVQ* deleted | | | This study |
| 1935 | 630∆*erm* ∆*sipL∆spoVQ*∆*pyrE* | | 630∆*erm* ∆*sipL∆pyrE* with *spoVQ* deleted | | | This study |
| 1962 | 630∆*erm ∆sleC*∆*spoVQ*-p | | 630∆*erm* ∆*sleC∆spoVQ* with *pyrE* restored | | | This study |
| 1965 | 630∆*erm ∆sipL*∆*spoVQ*-p | | 630∆*erm* ∆*sipL∆spoVQ* with *pyrE* restored | | | This study |
| 1968 | 630∆*erm ∆sipL*∆*spoVQ*/*sipL-mCherry* | | 630∆*erm* ∆*sipL∆spoVQ* with *sipL-mCherry* in the *pyrE* locus | | | This study |
| 1977 | 630∆*erm ∆cspC*∆*spoVQ*-p | | 630∆*erm* ∆*cspC∆spoVQ* with *pyrE* restored | | | This study |
| 1995 | 630∆*erm ∆cspC*∆*spoVQ*/*spoVQ* | | 630∆*erm* ∆*cspC∆spoVQ* with *spoVQ* in the *pyrE* locus | | | This study |
| 1998 | 630∆*erm ∆sleC*∆*spoVQ*/*spoVQ* | | 630∆*erm* ∆*sleC∆spoVQ* with *spoVQ* in the *pyrE* locus | | | This study |
| 2017 | 630∆*erm* ∆*spoIVA∆spoVQ*∆*pyrE* | | 630∆*erm* ∆*spoIVA∆pyrE* with *spoVQ* deleted | | |  |
| 2113 | 630∆*erm ∆spoIVA*∆*spoVQ*-p | | 630∆*erm* ∆*spoIVA∆spoVQ* with *pyrE* restored | | | This study |
| 2242 | 630∆*erm*∆*pyrE* ∆*cotL* | | 630∆*erm*∆*pyrE* with *cotL* deleted | | | This study |
| 2271 | 630∆*erm ∆cotL/cotL* | | 630∆*erm* ∆*cotL* with with *cotL* in the *pyrE* locus | | | This study |
| 2277 | 630∆*erm ∆cotL-p* | | 630∆*erm* ∆*cotL* with *pyrE* restored | | | This study |
| 2318 | 630∆*erm*/*spoVQ-mCherry* | | 630∆*erm* with *spoVQ-mCherry* in the *pyrE* locus | | | This study |
| 2321 | 630∆*erm ∆sipL*∆*spoVQ*/*spoVQ-mCherry* | | 630∆*erm* ∆*sipL∆spoVQ* with *spoVQ-mCherry* in the *pyrE* locus | | | This study |
| 2324 | 630∆*erm ∆spoIVA*∆*spoVQ*/*spoVQ-mCherry* | | 630∆*erm* ∆*spoIVA∆spoVQ* with *spoVQ-mCherry* in the *pyrE* locus | | | This study |
| 2442 | 630∆*erm ∆sipL*/*mCherry-sipL* | | 630∆*erm* ∆*sipL* with *mCherry-sipL* in the *pyrE* locus | | | This study |
| 2827 | 630∆*erm ∆sipL*∆*spoVQ*/*mCherry-sipL* | | 630∆*erm* ∆*sipL∆spoVQ* with *mCherry-sipL* in the *pyrE* locus | | | This study |
| ***E. coli* strains** | | |  | | |  |
| **Strain #** | **Strain name** | | **Relevant genotype or features** | | | **Source** |
| 41 | DH5α | | F– Φ80*lacZ*ΔM15 Δ(*lacZYA-argF*) U169 *recA1 endA1 hsdR17* (rK^–^, mK^+^) *phoA supE44* λ– *thi-1 gyrA96 relA1* | | | D. Cameron |
| 531 | HB101/pRK24 | | F- *mcrB mrr hsdS20*(rB^–^mB^–^) *recA13 leuB6 ara-13 proA2 lavYI galK2 xyl-6 mtl-1 rpsL20* carrying pRK24 | | | C. Ellermeier |
| 903 | BL21(DE3) pET28a + pET21a *sipL*(TAA) | | pET28a + pET21a *sipL*(TAA) | | |  |
| 1768 | HB101 pMTL-YN1C *mCherry-spoIVA* | | pMTL-YN1C *mCherry-spoIVA* | | | (9) |
| 1777 | HB101 pMTL-YN1C *sipL-mCherry* | | pMTL-YN1C *sipL-mCherry* in HB101 | | | (11) |
| 642 | DH5α pET21a-*spoIVA*(TAA) | | pET21a-*spoIVA*(TAA) | | | (2) |
| 643 | DH5α pET21a-*sipL*(TAA) | | pET21a-*sipL*(TAA) | | | (2) |
| 1975 | DH5α pET28a-*spoVQ*-His_6_ | | pET28a-*spoVQ*_∆32_-His_6_ | | | This study |
| 1976 | BL21(DE3) pET28a-*spoVQ*-His_6_ | | pET28a-*spoVQ*-His_6_ | | | This study |
| 1978 | BL21(DE3) pET28a-*spoVQ*_∆32_-His_6_ | | pET28a-*spoVQ*_∆32_-His_6_ | | | This study |
| 1996 | HB101 pMTL-YN3 ∆*spoVQ* | | pMTL-YN3 ∆*spoVQ* in HB101 | | | This study |
| 2050 | HB101 pMTL-YN1C *spoVQ* | | pMTL-YN1C *spoVQ* in HB101 | | | This study |
| 2052 | HB101 pMTL-YN1C *spoVQ-FLAG_3_* | | pMTL-YN1C *spoVQ-FLAG_3_* in HB101 | | | This study |
| 2054 | HB101 pMTL-YN1C *spoVQ-mCherry* | | pMTL-YN1C *spoVQ-mCherry* in HB101 | | | This study |
| 2056 | HB101 pMTL-YN1C *spoVQ*_∆32_ | | pMTL-YN1C *spoVQ*_∆32_ in HB101 | | | This study |
| 2355 | HB101 pMTL-YN1C *mCherry-sipL* | | pMTL-YN1C *mCherry-sipL* in HB101 | | | This study |
| 2067 | BL21(DE3) pET22b-CPD(TAA) | | pET22b-CPD(TAA) | | | This study |
| 2100 | BL21(DE3) pET28a-*spoVQ*_∆32_-His_6_ + pET21a | | pET28a-*spoVQ*_∆32_-His_6_ + pET21a | | | This study |
| 2057 | BL21(DE3) pET28a-*spoVQ*_∆32_-His_6_ + pET21a-*sipL*(TAA) | | pET28a-*spoVQ*_∆32_-His_6_ + pET21a-*sipL*(TAA) | | | This study |
| 2058 | BL21(DE3) pET28a-*spoVQ*_∆32_-His_6_ + pET21a-*spoIVA*(TAA) | | pET28a-*spoVQ*_∆32_-His_6_ + pET21a-*spoIVA*(TAA) | | | This study |
| 2071 | BL21(DE3) pET28a-*spoVQ*_∆32_-His_6_ + pET21a-CPD(TAA) | | pET28a-*spoVQ*_∆32_-His_6_ + pET21a-CPD(TAA) | | | This study |
| 2102 | BL21(DE3) pET28a + pET21a-CPD(TAA) | | pET28a + pET21a-CPD(TAA) | | | This study |
| 2239 | pMTL-YN3-∆*cotL* | | pMTL-YN3 ∆*cotL* in HB101 | | | This study |
| 2278 | pET22b-*cotL* | | pET22b-*cotL* in BL21(DE3) | | | This study |
| 2261 | pMTL-YN1C-*cotL* | | pMTL-YN1C *cotL* in HB101 | | | This study |
| 2354 | pMTL-YN1C *mCherry-sipL* | | pMTL-YN1C *mCherry-sipL in* | | | This study |

**Plasmids**

| pET28a | For cloning His-tagged expression constructs | Novagen |
| --- | --- | --- |
| pET21a | For cloning His-tagged expression constructs | Novagen |
| pMTL-YN1C | For cloning complementation constructs to be integrated into the pyrE locus of 630∆*erm*∆*pyrE* | (12) |
| pMTL-YN3 | For cloning allelic exchange constructs to modify 630∆*erm*∆*pyrE* | (12) |

**Table S2. Primers used in this study.**

| **Number** | **Primer Name** | **Primer Sequence** |
| --- | --- | --- |
| 2133 | 3' XhoI *mCherry* Gibson | gccaagcttgcatgtctgcaggcCTCGAGTTATTTATATAATTCATCCATACCTCCTGTTG |
| 2462 | 5' NcoI *spoVQ* | AGCCCATGGCATTGAATGTAAAGTTTAATATTAAAGGTATAATTTATG |
| 2463 | 3' XhoI *spoVQ* | GAGCCTCGAGTTTCGTTGCATTTAATTCTATTCTATCTGGAAG |
| 2464 | 5' NcoI *spoVQ*_∆32_ | AGCCCATGGCATTTGTAGAGAAATCTAAACCCATAGATTATACAG |
| 2484 | 5' AscI ∆*spoVQ* Gibson | gtcaattgttcaaaaaaataatggcGGCGCGCCTTATGGCATTCTTAACTGGATTAGGAG |
| 2485 | 5' ∆*spoVQ* SOE | GAGTTATAGCTCTTGTATTCGTTATAGGGGAACTTCCAGATAGAATAGAATTAAATGCA |
| 2486 | 3' ∆*spoVQ* rev eos | TGCATTTAATTCTATTCTATCTGGAAGTTCCCCTATAACGAATACAAGAGCTATAACTC |
| 2487 | 3' SbfI ∆*spoVQ* Gibson | gcaaggcaagaccgatcgggcccCCTGCAGGCTTCAAATGGAAGATATAAGAAATGCAAC |
| 2540 | 5' NotI *spoVQ* Gibson | ggaattagggatgtaataagcggccgcCAAATAAGTATTTTTTAATATTGTAAAC |
| 2541 | 3' *spoVQ* mCherry eos | CTTCTTCTCCTTTAGATACCATTGCTTTCGTTGCATTTAATTCTATTC |
| 2542 | 5' mCherry *spoVQ* SOE | GAATAGAATTAAATGCAACGAAAGCAATGGTATCTAAAGGAGAAGAAG |
| 2543 | 3' XhoI *spoVQ* Gibson | ttgcatgtctgcaggcctcgagTTATTTCGTTGCATTTAATTCTATTC |
| 2544 | 3' mid *spoVQ* for FLAG | CTTTCTATTTTGTCCATTTCTATACCATATTTATTATTTCCTCTAGTCAC |
| 2549 | 3' 5'UTR *spoVQ_∆32_* rev | CTATGGGTTTAGATTTCTCTACAAATGCCATTCTTATCACTCCTCCCATATATTC |
| 2550 | 5' YN1C *spoVQ_∆32_* SOE | GAATATATGGGAGGAGTGATAAGAATGGCATTTGTAGAGAAATCTAAACCCATAG |
| 2573 | 5' NdeI CPD 22b | TTTGTTTAACTTTAAGAAGGAGATATACATATGGCATTAGCGGATGGAAAAATACTCC |
| 2574 | 3' XhoI CPD+TAA 22b | CAGTGGTGGTGGTGGTGGTGCTCGAGTTAACCTTGCGCGTCCCAGCTTAGCGAAAC |
| 2820 | 5' AscI ∆*cotL* Gibson | gtcaattgttcaaaaaaataatggcggcgcgccGAGATTATTCATACTACCAAGATTG |
| 2821 | 5' ∆*cotL* SOE | TGAATATAATTCAATATATATTTATAGAAAGGGGTAAAACTAAAAAGTCTAACCCTCATG |
| 2822 | 3' ∆*cotL* rev eos | CATGAGGGTTAGACTTTTTAGTTTTACCCCTTTCTATAAATATATATTGAATTATATTCA |
| 2823 | 3' SbfI ∆*cotL* Gibson | agcaaggcaagaccgatcgggccccctgcaggGAGTATAAATCTATTGAGAGC |
| 2833 | 5' NotI *cotL* Gibson | aattagggatgtaataagcggccgcCTTAAATTGCCATGTGTAGACTGTG |
| 2834 | 3' XhoI *cotL* Gibson | caagcttgcatgtctgcaggcctcgagCTATTCATGAGGGTTAGACTTTTTAG |
| 2946 | 5’ NdeI *cotL* pET22b | TTTAACTTTAAGAAGGAGATATACATATGCTTAAATTGCCATGTGTAGACTGTGAAAG |
| 2947 | 3’ XhoI *cotL* pET22b | ATCTCAGTGGTGGTGGTGGTGGTGCTCGAGTTCATGAGGGTTAGACTTTTTAGTTTTC |
| 3019 | 5’ *mCherry-sipL* SOE | CAGGAGGTATGGATGAATTATATAAAGCAATGGAATTAATTAAAGATGTAATTAAAG |
| 3020 | 3’ *mCherry-sipL* EOS | CTTTAATTACATCTTTAATTAATTCCATTGCTTTATATAATTCATCCATACCTCCTG |
| 3021 | 5’ P*sipL*-*mCherry* SOE | ATATTTTTATAATACTTAAGGAGGTAGACTATGGTATCTAAAGGAGAAGAAGATAATATG |
| 3022 | 3’ P*sipL*-*mCherry* EOS | CATATTATCTTCTTCTCCTTTAGATACCATAGTCTACCTCCTTAAGTATTATAAAAATAT |

Restriction sites are underlined.
